# A protein-RNA interaction atlas of the ribosome biogenesis factor AATF

**DOI:** 10.1101/446260

**Authors:** Rainer W. J. Kaiser, Michael Ignarski, Eric L. Van Nostrand, Christian Frese, Manaswita Jain, Sadrija Cukoski, Heide Heinen, Melanie Schaechter, Lisa Seufert, Konstantin Bunte, Peter Frommolt, Patrick Keller, Mark Helm, Katrin Bohl, Martin Höhne, Bernhard Schermer, Thomas Benzing, Katja Höpker, Christoph Dieterich, Gene W. Yeo, Roman-Ulrich Müller, Francesca Fabretti

## Abstract

AATF is a central regulator of the cellular outcome upon p53 activation, a finding that has primarily been attributed to its function as a transcription factor. Recent data showed that AATF is essential for ribosome biogenesis and plays a role in rRNA maturation. AATF has been implicated to fulfil this role through direct interaction with rRNA and was identified in several RNA-interactome capture experiments. Here, we provide a first comprehensive analysis of the RNA bound by AATF using CLIP-sequencing. Interestingly, this approach shows predominant binding of the 45S pre-ribosomal RNA precursor molecules. Furthermore, AATF binds to mRNAs encoding for ribosome biogenesis factors as well as snoRNAs. These findings are complemented by an in-depth analysis of the protein interactome of AATF containing a large set of proteins known to play a role in rRNA maturation with an emphasis on the protein-RNA-complexes known to be required for the generation of the small ribosomal subunit (SSU). In line with this finding, the binding sites of AATF within the 45S rRNA precursor localize in close proximity to the SSU cleavage sites. Consequently, our multilayer analysis of the protein-RNA interactome of AATF reveals this protein to be an important hub for protein and RNA interactions involved in ribosome biogenesis.

## INTRODUCTION

The apoptosis antagonizing transcription factor (AATF), also known as Che-1, was originally identified as an RNA polymerase II interacting protein with anti-apoptotic capacities. Through its interaction with RNAPII, AATF modulates the function of a row of transcription factors including pRB, p65 and STAT3, and has been shown to be a transcription factor itself^1–3^. Considering its role in the prevention of apoptosis, the involvement of AATF in the DNA damage response and its impact on p53 function are explicitly interesting. The functional impact of these findings is emphasized by data showing AATF as a pro-tumorigenic factor in several tumor models^4,5^. Upon DNA damage AATF is phosphorylated by checkpoint kinases leading to its interaction with the NF-kB subunit p65 and relocalization to the p53 promoter^3,6,7^. However, AATF also modulates the specificity of p53 by shifting its binding preference towards target genes leading to growth arrest over those that mediate apoptosis^5,8^. Furthermore, AATF has been shown to play a role in other key pathways involved in tumorigenesis, namely mTOR- and HIF-signaling^9,10^. Seeing the impact of this protein on signal transduction in apoptosis and tumor formation, its potential as a therapeutic target in cancer therapy has been discussed extensively^5,11,12^. Interestingly, AATF – whilst also detected in the cyto- and nucleoplasm – primarily localizes to the nucleolus with the nucleolar fraction mediating the impact of AATF on c-JUN-dependent apoptosis^13^. The strongly regulated biogenesis of ribosomes starts in the nucleolus where transcription and most of the processing of the large primary transcript (45S rRNA) occur. This 45S rRNA contains the 18S, 5.8S and 28S rRNAs interspersed by internal and external transcribed spacers (ITS1/2 and 5’-/3’-ETS). To create the immense molecular machine of a ribosome, ribosomal proteins (RPs), pre-ribosomal factors (PRFs) and small nucleolar RNAs (snoRNAs) assemble with the 45S rRNA to support pre-RNA folding, deposit RNA modifications and to remove ITS and ETS regions by endonucleolytic and exonucleolytic cleavage as reviewed by Henras et al.^14^. Ultimately, in mammalian cells the 60S large subunit is assembled from the 5S, 5.8S, 28S rRNAs and 46 ribosomal proteins (RPs) whereas the 40S small subunit contains the 18S rRNA and 33 RPs. It is important to note that AATF was recently identified by independent RNAi screens for factors involved in ribosomal subunit production and rRNA processing^15,16^. This connection is interesting due to two considerations. Firstly, cell proliferation and ribosome biogenesis rate are closely intertwined, and increased ribosome formation is linked to tumorigenesis^17,18^. Secondly, blocking ribosome formation induces ribosome biogenesis stress leading to the activation of p53 mediated by inhibition of MDM2. Vice versa, induction of ribosome biogenesis inhibits p53^17,19,20^. However, the mode of action of AATF in ribosome maturation has remained unclear. A first insight came recently from a study by Bammert et al.^21^ that could show AATF as part of a nucleolar protein complex (termed ANN complex). Here, AATF, together with NOL10 and NGDN, was essential to efficient generation of the small ribosomal subunit (SSU) - in line with findings from previous screens of ribosome biogenesis factors^15^. Yet, the molecular function that AATF fulfils within this complex and the question which one of the three proteins mediates binding to rRNA precursors remained elusive. Work published recently shows AATF to be amongst 211 RNA polymerase I-dependent RNA-binding proteins^22^. However, a global characterization of its RNA interaction partners and binding sites that allows for a better understanding of its functional implications in ribosome biogenesis were missing. We confirmed AATF to be an RNA-binding protein and extended the analysis to a comprehensive characterization of its RNA-interactome using RNA-sequencing of its targets after crosslinking and immunoprecipitation (eCLIP-seq). Our combination of this approach – showing AATF to bind primarily rRNA precursor molecules and mRNAs encoding proteins required for ribosome biogenesis – with the analysis of its protein interactome suggests AATF to be a central player in the coordination of protein and RNA components in the maturation of the mammalian SSU.

## RESULTS

### AATF is an RNA-binding protein associated with ribosomal RNA

Previous studies from our own group using RNA interactome capture (RIC) of cultured mouse inner medullary collecting duct (mIMCD3) cells identified the highly conserved mouse orthologue of AATF, Aatf/Traube, as one of the most enriched proteins in the RNA-bound proteome (Suppl. Fig. 1A)^23^. In order to confirm this finding and to show it is not limited to our experiment we screened for the presence of AATF in published RIC datasets. Despite the fact that these studies examined different species and cell lines and used different experimental strategies, AATF as well as its mouse (Aatf/Traube) and yeast (Bfr2) orthologues were among the most strongly enriched proteins (Suppl. Fig. 1A)^24–29^. In order to confirm its binding to RNA and to determine the identity of the RNA molecules bound to AATF, we analyzed enhanced crosslinking and immunoprecipitation (eCLIP) data generated by the ENCODE consortium^30^ (Suppl. Fig. 1B). The majority of transcripts bound to AATF are ribosomal RNAs (Fig. 1A). This finding is of interest considering the subcellular localization of AATF to the nucleolus, the site of rRNA transcription, initial cleavage and modification of pre-rRNA, and considering as well that previously published data showed rRNA maturation defects upon loss of AATF^16,21,22,31^,. Despite rRNA being the most abundant RNA biotype, AATF shows a much higher signal for rRNA than both the input controls and the eCLIP results of −223 other publicly available datasets of 150 RNA binding proteins (RBPs) generated by the ENCODE consortium (Fig. 1A). Whilst – as expected – the nucleolar RBPs show more rRNA binding in general than non-nucleolar RBPs, the median fraction of rRNA species bound is lower than for AATF with many nucleolar RBPs not binding to rRNA (Fig. 1A). The overrepresentation is strongest for the 45S rRNA (Fig. 1B), the early precursor of mature 28S, 18S and 5.8S ribosomal RNA that is transcribed from rDNA in the nucleolus. In line with its role in rRNA biogenesis, AATF has been described as being primarily nuclear with several reports confirming an accumulation of the protein in nucleoli^13,21^. Previous work indicated that the C-terminal portion of AATF is required for its specific nucleolar localization. However, the actual nucleolar localization signals (NoLS) had not been identified^13,21^. The NucleOlar localization sequence Detector (NoD)^32^ predicts two putative NoLS in the C-terminal portion of the protein (amino acid position 326 to 345 and position 494 to 522) (Suppl. Fig. 1C). In order to visualize the subcellular localization of AATF, we generated transgenic U2OS cell lines stably expressing a single-copy transgene encoding GFP-tagged versions of either WT AATF or a mutant protein lacking the two NoLS stretches, (AATF 2ΔNoLS truncation, Suppl. Fig. 1D). The WT fusion protein localizes predominantly to nucleoli, whilst the mutant protein is dispersed throughout the cyto- and nucleoplasm, confirming that the predicted sequences indeed serve as NoLS (Suppl. Fig. 1E). We then used overexpressed FLAG-tagged AATF (WT and 2ΔNoLS) to confirm specificity of the binding of AATF to 45S pre-rRNA and examine the impact of nucleolar localization on RNA binding. RNA immunoprecipitation coupled with quantitative PCR (RIP-qPCR) on overexpressed FLAG-tagged AATF showed a clear enrichment of rRNA over a control pulldown (FLAG tagged RFP, red fluorescent protein) with the most distinct signal for the 45S rRNA precursor (Fig. 1C). Of note, both wild-type AATF and the AATF 2ΔNoLS truncation were expressed at the same level (Suppl. Fig. 1F). As expected, loss of nucleolar localization of AATF resulted in a decreased association with rRNAs, in particular in case of the 45S rRNA precursor (Fig. 1C). Furthermore, knockdown of AATF lowered the cellular content of mature ribosomal RNA as shown by both EtBr staining (Fig. 1D) and RT-qPCR of 18S rRNA (Suppl. Fig. 1G). This loss of rRNA can be rescued using a single-copy transgene of AATF that lacks the 3’ untranslated region, i.e. binding site of the siRNA, confirming specificity of this finding (Fig. 1E).

**Figure 1:**
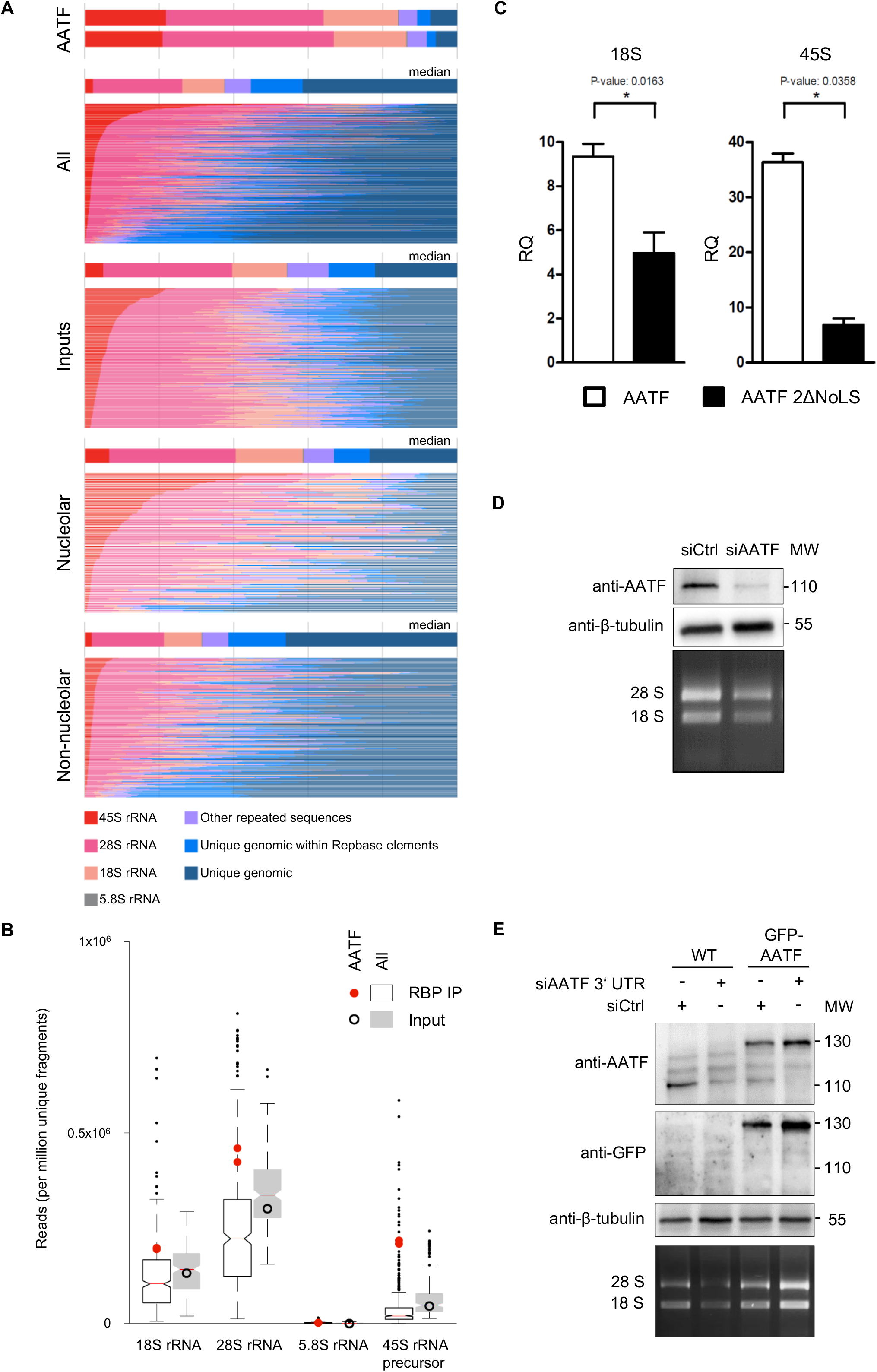
Analysis of AATF-bound RNA biotypes identified by eCLIP. **A** Stacked bar colors indicate the fraction of reads mapping to indicated ribosomal RNA, other repeat elements, or uniquely to the human genome. For comparisons the total number of ENCODE datasets were used (223 datasets of 150 RBPs)^59^. All = all datasets, inputs = all input controls, nucleolar = all nucleolar RBPs among the ENCODE datasets, non-nucleolar = all non-nucleolar RBPs among the ENCODE datasets; “unique genomic within Repbase elements” = mapping uniquely to a repeat-masker identified element in the genome, “other repeated sequences” = mapping to the canonical repetitive elements in the family analysis. Nucleolar or non-nucleolar localization of RBPs was based on immunofluorescence data from the ENCODE consortium (see methods section for details). **B** Box plot comparison of AATF eCLIP-seq data to input and to 150 ENCODE RBPs showing enrichment of rRNA species in IP (red dot) over input (black circle) is specific for AATF. The other 150 ENCODE RBPs show a decrease of rRNA species in the IP (white box) compared to inputs (grey box). RNA species are plotted against the reads identified per million unique fragments. **C** RIP-qPCR analysis of 18S rRNA and 45S pre-rRNA transcripts validating the capacity of AATF to bind rRNA. Full-length FLAG-tagged AATF, the FLAG-tagged AATF 2ΔNoLS truncation or FLAG-tagged RFP (red fluorescent protein) were transiently overexpressed in HEK 293T cells and immunoprecipitated in RNA-interaction preserving conditions (n = 3). Quantification of co-precipitated rRNA revealed a significant reduction of RNA binding for both ribosomal transcripts after loss of the two NoLS sites. RQ: relative quantification (CT values for WT AATF, 2ΔNoLS AATF and RFP were normalised against the corresponding input (delta CT_IP-INPUT_), and consecutively against RFP (delta delta CT, e.g. delta CT_AATF_ − delta CT_RFP_). FLAG-RFP served as negative control. For western blot of IP from whole cell lysates showing equal protein amounts see Suppl. Fig. 1F. **D** Knockdown of AATF leads to a reduction of rRNA. The CDS of AATF was targeted with siRNA in mIMCD3 cells, which induced a significant depletion of endogenous AATF and was accompanied by a decrease in rRNA after 48h of incubation. Top panel: western blot with anti-AATF antibody. Middle panel: anti-β-tubulin western blot (loading control). Bottom panel: EtBr stained agarose gel. MW: protein molecular weight marker (kDa). **E** Expression of AATF single-copy transgene rescues reduction of rRNA in AATF depleted U2OS cells. siRNA against the 3’UTR of AATF was transfected into wild-type U2OS cells and U2OS cells with a TALEN mediated, single-copy integration of GFP-AATF lacking the endogenous 3’UTR into the AAV locus. The 3’UTR specific knockdown of AATF in the wild type cells lead to a reduction of the 18S and 28S rRNA. The expression of the GFP-tagged transgene in the TALEN manufactured U2OS GFP-:AATF cell line rescued the amount the rRNA species. Top panel: western blot with anti-AATF antibody. Middle panel: anti-β-tubulin western blot (loading control). Bottom panel: EtBr stained agarose gel. MW: protein molecular weight marker (kDa).

### Specific binding of AATF to SSU cleavage sites in ribosomal precursor RNAs

Considering the described binding of AATF to 45S pre-rRNA, we focused our study on this interaction. We discovered a number of binding sites depicted by specific peaks along the 45S transcript (Fig. 2A). Compared to the 150 ENCODE RBPs, AATF peaks are strongly enriched when looking at the spacer regions, whilst less prominent regarding the 18S and 28S region. We therefore concentrated on the spacer sequences outside 18S and 28S rRNA. These spacer regions (5’ and 3’ external transcribed spacers (ETS) and internal transcribed spacers 1 and 2 (ITS)) are part of the 45S pre-rRNA transcript and are located outside (ETS) or in between the sequences encoding three ribosomal RNAs (ITS, Fig. 2A and 2B)^33^. Among a variety of processing steps such as RNA modification, nucleolar rRNA maturation involves endonucleolytic cleavage at specific sites in these spacer regions before export to the cytoplasm, where additional exonucleolytic maturation of 18S and 5.8S takes place before the final ribosome is assembled^14^. When examining the eCLIP peaks in the spacer regions in more detail, we noticed a close proximity to some of the endonucleolytic cleavage sites (Fig. 2B)^15^. This is especially the case for sites involved in small subunit (SSU) processing as depicted for the sites in the 5’ external transcribed spacer (5’ ETS) – 01, A0 and 1 – and the first sites in internal transcribed spacer 1 (ITS1) following the sequence of 18S (Fig. 2B)^16,34^. This finding is in line with recently published data showing that loss of AATF or other components of the so-called ANN complex results in reduced cleavage activity at these sites indicating reduced SSU processome activity^15,21^. When comparing these results to the other available ENCODE eCLIP datasets, peaks in the 45S pre-rRNA are in general only discovered for nucleolar RBPs (Suppl. Fig. 2). However, even among nucleolar RBPs only a subset shows specific peaks comparable to the ones found for AATF (Suppl. Fig. 2). Other non-SSU associated sites in ITS1, ITS2 and 3’ETS are also detected in the AATF eCLIP but show a much lower signal (e.g. site 2, Fig. 2B).

**Figure 2:**
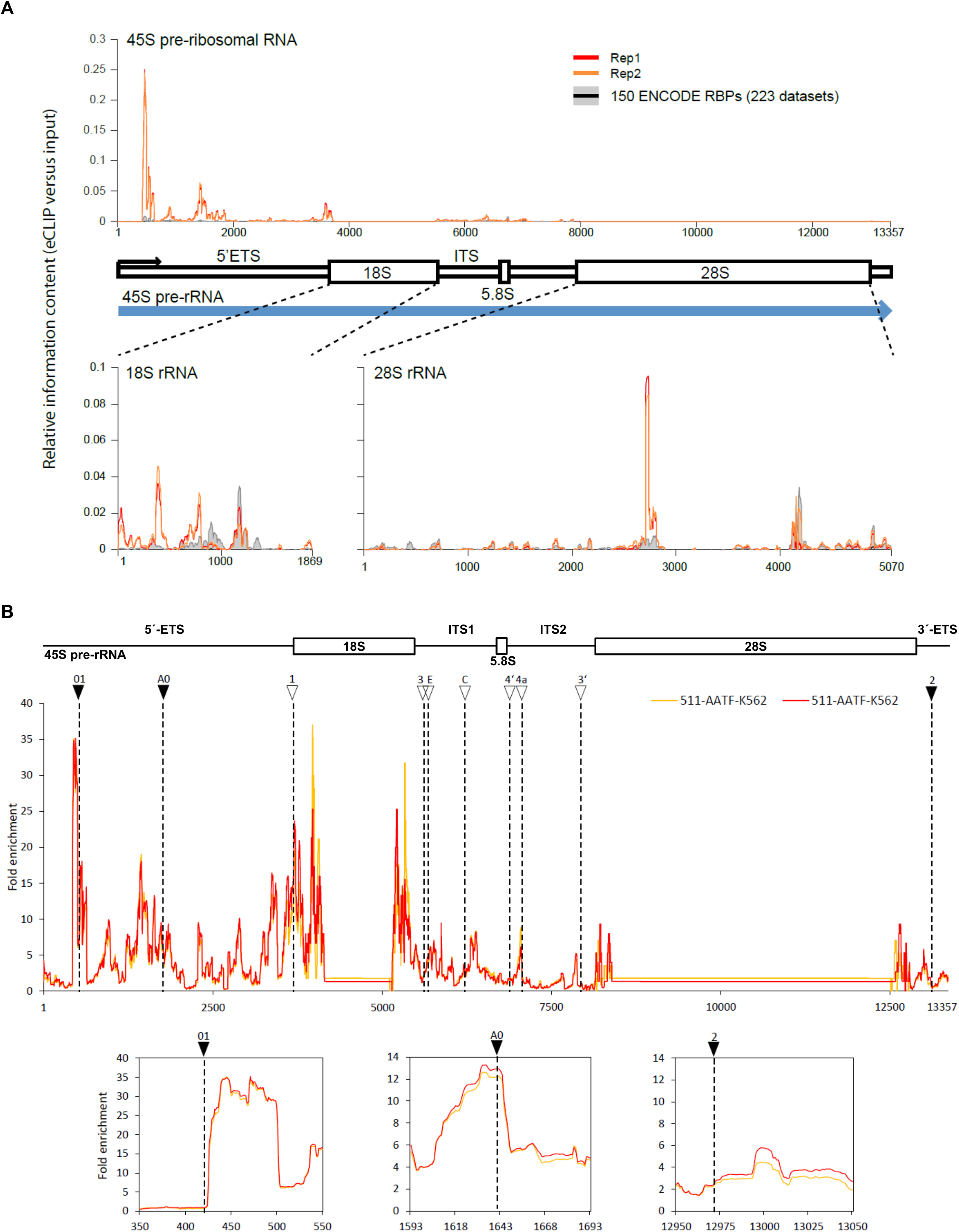
Identification of 45S pre-rRNA sites bound by AATF. **A** Relative information content on eCLIPseq peak distribution along the 45 pre-rRNA and the mature 18S and 28S rRNA comparing AATF to all ENCODE RBP datasets. **B** AATF eCLIP reads map to the 45S pre-rRNA and show enrichment at cleavage sites involved in SSU maturation. Top: Scheme of 45S pre-rRNA. Middle: Graph showing the fold enrichment of eCLIP reads of two replicates (yellow and red) along the 13357 bp long 45S pre-rRNA. 45S rRNA cleavage sites involved in SSU maturation are indicated by arrows and dashed lines. Bottom: Black arrow heads indicate regions shown below in detail for the sites: 01, A0 and 2.

### AATF binding to other RNA species

As indicated in Fig. 1A, AATF binds also to non rRNA-species of which mRNA is the most common type (69% protein coding) followed by several types of non-coding RNAs (Fig. 3A, Suppl. Table 1). Seeing the association with both ribosomal RNA and mRNAs we were wondering whether the associated coding transcripts were functionally linked to ribosome biogenesis. Indeed, gene ontology (GO) and pathway enrichment analyses confirmed such a link with the most overrepresented terms including “rRNA processing”, “ribosome”, “translational initiation” and “structural constituent of ribosome” regarding gene ontologies and “ribosome” for KEGG pathways (Fig. 3B). As shown in Fig. 3A several non-coding RNA-species interact with AATF including snoRNAs. This finding is especially interesting taking into account the important role of snoRNAs and associated snoRNPs in ribosome biogenesis^17,35,36^. Interestingly, snoRNAs of the C/D class are more commonly detected in the AATF eCLIP (n=69 out of 95) compared to H/ACA snoRNAs (n=19) and scaRNAs (n=7) indicating a specific function associated with C/D box snoRNPs (Fig. 3C)^37^. Indeed, AATF is among the 5 RBPs (compared to total ENCODE datasets) with the strongest fold enrichment of C/D box snoRNAs and no such overrepresentation is detected for H/ACA box snoRNAs (Fig. 3D). Since C/D box snoRNAs are key players in site-specific nucleotide modification of ribosomal RNA precursors and play an important role in their maturation, we analysed whether loss of AATF may have a global impact on the abundance of these modifications. However, quantification of the most common rRNA modifications by mass spectrometry did not reveal any significant differences between cells transfected with an siRNA targeting AATF and the respective controls (Suppl. Fig. 3).

**Figure 3:**
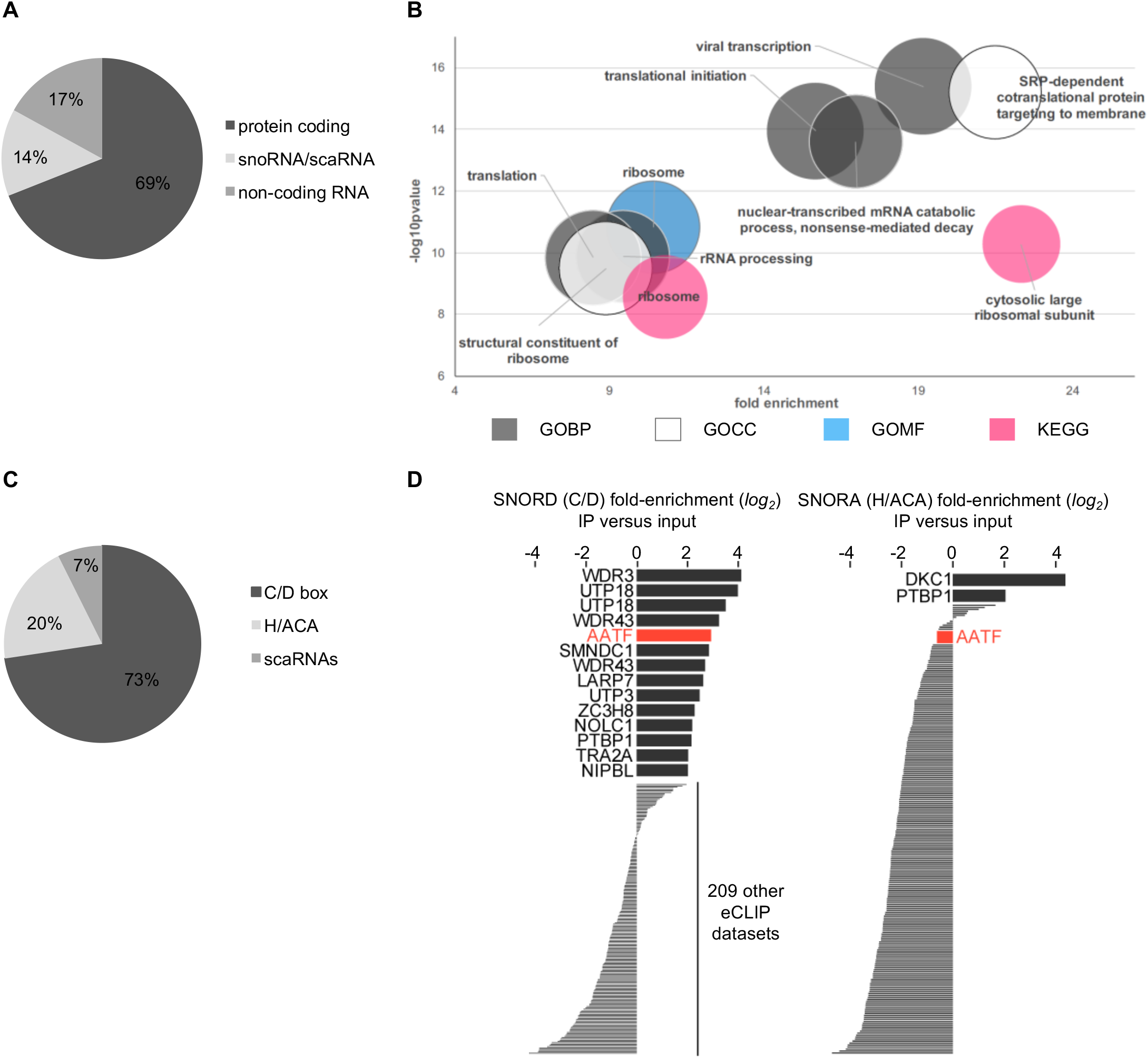
AATF interacts with coding and non-coding RNA species. **A** Pie chart depicting the distribution of RNA biotypes bound by AATF other than rRNA. 69% of transcripts other than rRNA bound by AATF are protein-coding, (14%) snoRNA and scaRNA, and “non-coding RNA” biotypes (17%) encompass lincRNA, miRNA and antisense RNA. **B** Bubble chart depicting the functional analysis of mRNA transcripts bound by AATF showing the terms contained in the top functional annotation cluster as identified using the DAVID Bioinformatics online tool^67^ (for the 292 eCLIP targets showing significant peaks in at least two experiments, Suppl. Table 1). GO terms are plotted according to fold enrichment and −log_10_ of the respective p-value, with size of the bubble increasing proportionally the number of genes contained in the respective cluster. **C** Pie chart showing the proportions of snoRNAs bound by AATF. Among the transcript biotype group of snoRNAs AATF preferentially binds C/D box snoRNAs, with 73% of bound snoRNAs belonging to this subtype. Box H/ACA snoRNAs comprise 20% and scaRNA 7% of transcripts bound by AATF. **D** Bars indicate the fold-enrichment (or depletion) in immunoprecipitation versus input for C/D-box snoRNAs and H/ACA-box snoRNAs in all ENCODE eCLIP datasets. AATF is noted in red, and other datasets with at least 4-fold enrichment are indicated by name.

### The AATF protein interactome confirms a strong link to ribosome biogenesis

Since AATF had been associated with other nucleolar proteins before ^21^ and the recruitment of components of the ribosome biogenesis machinery to the 45S precursor may be one of the key functions of an rRNA-associated RBP, we decided to characterize the protein interactome of AATF by affinity-purification mass spectrometry (AP-MS) of FLAG-tagged AATF expressed from a single-copy transgene. In parallel, an immunoprecipitation in a control cell line expressing FLAG-tagged GFP to compare the abundance of proteins in both datasets was performed. We identified 165 proteins significantly enriched by AATF precipitation (log_2_FC ≥2 and −log_10_pvalue ≥ 1.3) that had partly been discovered and validated before to be associated with AATF in yeast or ≥ mammals (Fig. 4A/Suppl. Table 2). To validate our dataset, we went on to confirm the interaction with three of these proteins known to play a role in SSU maturation using endogenous co-immunoprecipitations (Co-IP). Indeed, all three proteins – the rRNA 2’-O-methyltransferase fibrillarin (FBL), the rRNA methyltransferase Nucleolar Protein 2 Homolog (NOP2) and the ribosome biogenesis factor HEAT Repeat-Containing Protein 1 (HEATR1) – co-precipitated with AATF with no signal in IgG-only control precipitates (Fig. 4B). Interestingly, an analysis of the GO terms and KEGG pathways most overrepresented among all AATF protein interactors showed primarily terms associated with the ribosome and its biogenesis (Fig. 4C). A closer look at the proteins behind these terms revealed the interactome to contain a large number of known ribosomal proteins (r-proteins) and rRNA processing factors (as recently identified by several screens)^15,17^ as well as nucleic acid associated enzymes including helicases known to be involved in snoRNA binding or release such as DHX15^38^ (Fig. 4D). Interestingly, more than 80% of the bona fide interactors have been identified as RBPs in independent screens themselves (Suppl. Fig. 4A). Since the AATF interactome contains such a large number of putative or confirmed RBPs we were wondering whether these interactions were actually RNA-dependent, as had been shown previously for other RBPs^39^. However, repeating the AP-MS experiment including an on-column RNase/benzonase treatment showed that the majority of the interactors do not depend on RNA but are rather direct protein-protein interactions (Fig. 4E, Suppl. Table 2). Using stringent thresholds (log_2_FC ≥2, −log_10_pvalue ≥ 1.3 in the t-test performed between AATF IP treated with RNase versus GFP IP) 93 out of 165 interactors identified without RNase still reached our criteria in the experiment after RNase treatment (and 147 out of 165 did so for alleviated thresholds log_2_FC ≥1, −log_10_pvalue ≥ 1.0) (Suppl. Table 2). This does not necessarily mean that the other 72 (18) proteins are RNA-dependent interactors: When comparing the +/− RNase AATF pulldowns directly to each other only two proteins reached stringent thresholds for being overrepresented when RNA is present (log_2_FC ≤−2, −log_10_pvalue ≥1.3 in the t-test performed between AATF IP treated with RNase versus AATF IP normalized) (Fig. 4E, Suppl. Table 2). In order to obtain a better view of which proteins may at least show a partial dependency on RNA we repeated this analysis using alleviated thresholds (log_2_FC ≤−1, −log_10_pvalue ≥1.0) which still results in only 19 putatively RNA-dependent interactors of AATF (Fig. 4E, Suppl. Table 2). Interestingly, a significant proportion of these 19 proteins (~30%) are classical ribosomal proteins, whilst this is only the case for 21 out of 146 of the other AATF interactors (14%) (Suppl. Fig. 4B). However, GO-term and KEGG-pathway enrichment analyses did not reveal any further differences between putatively RNA-dependent and -independent protein interactors (data not shown). Only two interactors were completely lost after nuclease treatment – RNA-binding protein SLIRP and protein phosphatase PPP1CB (Fig. 4E [indicated by red arrows], Suppl. Table 2). Next, since AATF has recently been shown to be RNAPI-associated, we asked whether the large number of RBPs in its interactome contained other RNAPI-dependent RBPs. Interestingly, the overlap of 105 proteins between our AATF interactome data with the recent global identification of RNAPI-dependent RBPs by Piñeiro et al.^22^ revealed AATF to interact with both RNAPI-dependent (46) and -independent (59) RBPs (Suppl. Fig. 4C/D).

**Figure 4:**
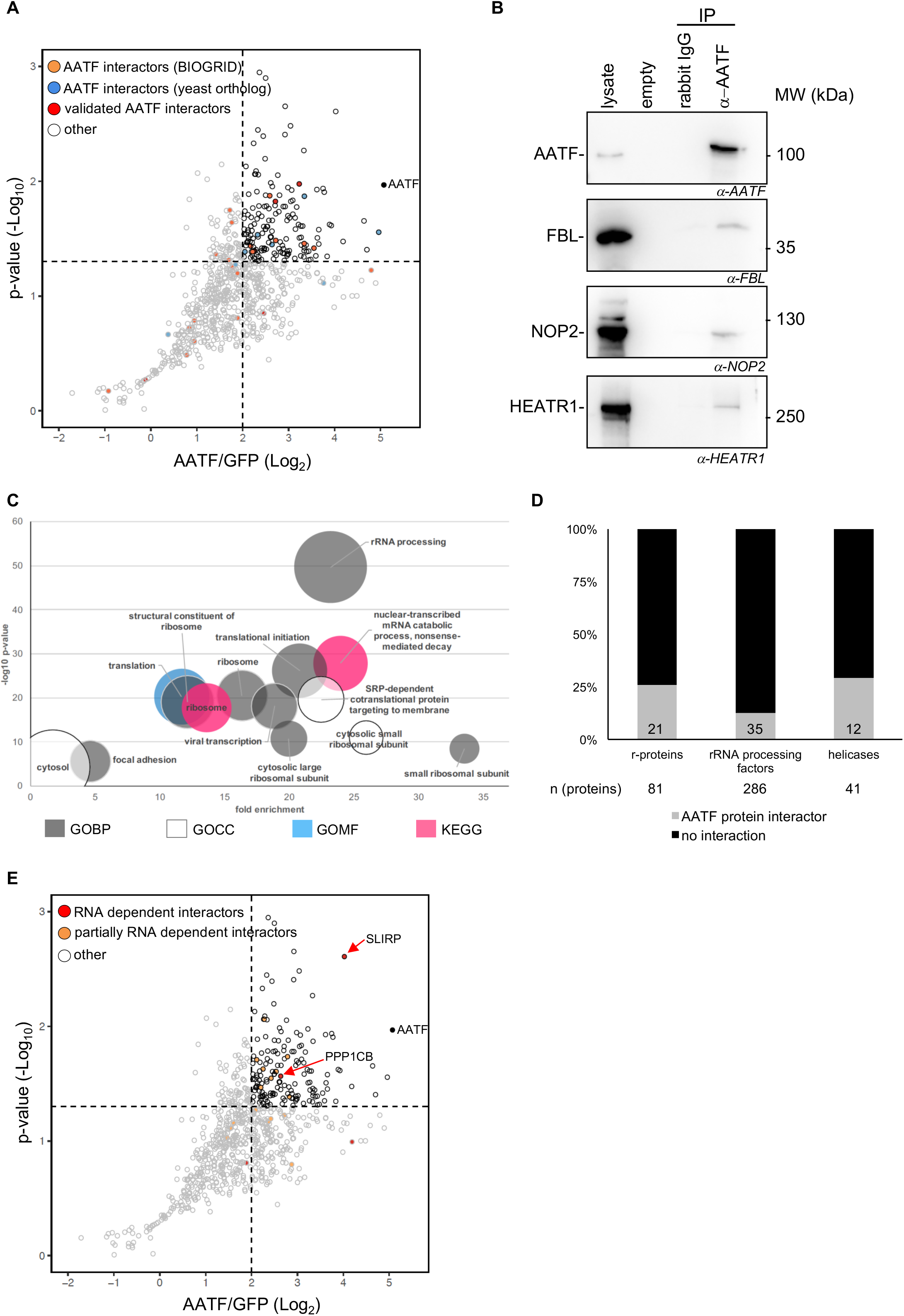
The protein interactome of AATF is strongly enriched for proteins involved in ribosome biogenesis. **A** Scatter plot of AATF interactome. Immunoprecipitation of FLAG-AATF (expressed from single-copy insertion in FlpIn 293T cells) and mass spectrometric analysis revealed 165 protein interactors fulfilling stringent criteria when compared to a FLAG-GFP pulldown (log_2_FC ≥2, −log_10_ p-value ≥1.3, n = 5). Known interactors are labeled with colored dots (black dot: AATF, orange dots: physical interaction as annotated in BIOGRID for human AATF, blue dots: physical interaction as annotated for the yeast AATF ortholog, red dots: AATF interactors experimentally validated in literature, white dots: previously not identified interactors). **B** Co-immunoprecipitation of endogenous AATF and western blot of novel interactors: 3 representative novel interactors identified in MS experiments that are known to be involved in SSU maturation (FBL, HEATR1 and Nop2/NSUN1) were confirmed by co-IP and western blot (n=3). Rabbit IgG only was used as negative control. MW = molecular weight marker (kDa). **C** Bubble chart depicting the functional analysis of AATF interacting proteins showing the terms contained in the top functional annotation cluster as identified using the DAVID Bioinformatics online tool^67^ (for the 165 bona fide AATF interactors). GO terms are plotted according to fold enrichment and −log_10_ of the respective p-value, with size of the bubble increasing proportionally the number of genes contained in the respective cluster. **D** Bar chart showing the percentage of AATF interacting proteins in the protein groups of r-proteins^17^, rRNA processing factors^15^ and human RNA helicases^27^ in grey. The numbers below indicate the total number of proteins per group, the numbers within the grey bars indicate the number of AATF interactors within this group. **E** Scatter plot highlighting RNA dependent AATF interacting proteins. Comparing the interactome after RNase treatment revealed that only few of the protein interactions depend on RNA (black dot: AATF, red dots: RNA-dependent interactions as defined by a log_2_FC ≥2 and −log_10_ p-value ≥1.3 compared to RNase treated IP, orange dots: partially RNA-dependent interactions as defined by a log_2_FC ≥1 and −log_10_ p-value ≥1 compared to RNase treated IP, white dots: RNA independent interactors). See also Suppl. Figure 4D for a direct comparison and Suppl. Table 2.

### A combinatorial approach to the RNA and protein interactome suggests AATF to be a key player in the coordination of the protein-RNA supercomplexes in SSU maturation

Considering the broad interaction with protein and RNA components of the SSU maturation complexes and previously published data suggesting the yeast AATF orthologue Bfr2 to be part of the SSU processome^40,41^ we focused our analysis of the AATF protein and RNA interactome on the key complexes involved in SSU maturation (Fig. 5A-F). Here, AATF shows a striking interaction with protein components of the three UTP complexes - tUTP, UTP-B and UTP-C (Fig. 5A-C). Among constituents of the tUTP complex, which is essential for rRNA transcription and association with pre-18S rRNA^34^, AATF shows a stringent interaction with HEATR1 (also confirmed by Co-IP, Fig. 4B) and WDR43 at protein level, while three of the four remaining tUTP constituents are also associated with AATF on either RNA and/or protein level (Fig. 5A). Looking at the components of the UTP-B complex, an SSU sub-complex required for rRNA 2’-O-methylation^42^, we found that AATF binds seven out of the eight UTP-B proteins (including NOP2 confirmed by Co-IP, Fig. 4B) and mRNAs encoding 5 of the respective proteins. Only for PWP2 neither an interaction at RNA nor at protein level was detectable (Fig. 5B). All five components of the UTP-C complex, which is thought to phosphorylate and thereby regulate other SSU complexes^34,41^, are bound by AATF at either RNA or protein level when applying alleviated thresholds (Fig. 5C). Furthermore, AATF does not only bind the proteins but also several mRNAs of the C/D snoRNP and MPP10 complex, containing essential rRNA methylation factors such as FBL (also confirmed by Co-IP, Fig. 4B) and multiple snoRNAs. Analysis of this network showed binding of most protein components. Furthermore, our eCLIP-seq data showed AATF to be associated with 69 C/D box snoRNAs (Fig. 3C) including U3 and U8 (Fig. 5D and Suppl. Table 1). As to the H/ACA snoRNP, our data primarily support an interaction on the transcript level, including the enzymatic subunit DKC1 that mediates the conversion of uridine to pseudouridine^34,41^. Besides, CLIP-sequencing identified 19 H/ACA snoRNAs to be co-precipitated with AATF (Fig. 5E). Finally, our data revealed an interaction with the exosome complex, a protein complex with exoribonuclease activity involved in the processing of numerous RNA biotypes including rRNA^34,41^. 9 of its 11 components are detected in CLIP-sequencing and/or by mass spectrometry after immunoprecipitation of AATF including the enzymatic subunits DIS3 and RRP6 (Fig. 5F).

**Figure 5:**
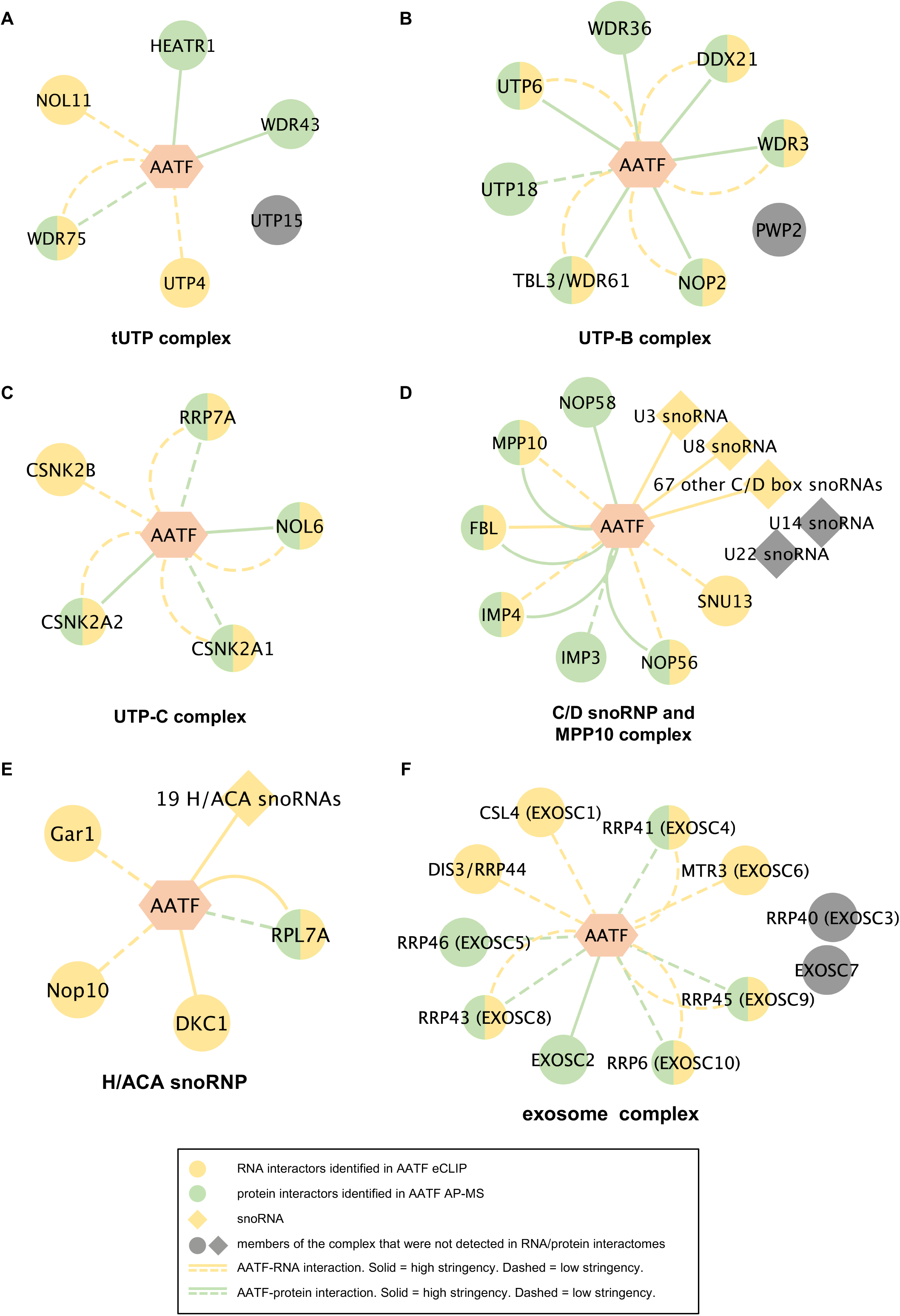
AATF is a central component of the protein complexes involved in SSU maturation and interacts with RNAs and proteins involved. **A-F** In our dataset, AATF interacts with the majority of the proteins known to be members of the key protein complexes involved in the maturation of ribosomal RNA^34,41^. Networks were created using Cytoscape and show constituents of the tUTP (A), the UTP-B (B), the UTP-C (C), the C/D snoRNAP and MPP10 complex (D) as well as the H/A snoRNP (E) and the exosome complex (F). (stringency-eCLIP (see Suppl. Table 1): high = 665 eCLIP targets containing at least one significant peak, low = all targets identified in eCLIP; stringency AP-MS: high = log_2_FC ≥ 2, log_10_ p-value ≥ 1.3 in either AATF with or without RNAse, low = detected in our interactome with a positive FC in at least two replicates of AATF with or without RNAse, both compared to the respective GFP pulldown)

## DISCUSSION

AATF had previously been associated with ribosome biogenesis due to its nucleolar localization and the impact of loss of the protein on the generation of the 40S subunit and rRNA maturation^15,16^. In our study, depletion of AATF led to a reduction in mature rRNA levels and previously published work had shown an accumulation of rRNA precursors upon loss of AATF^15,21^. Furthermore, AATF has been shown to be part of a protein complex (ANN complex) required for efficient generation of the SSU^21^. However, the actual RBP associating this protein complex to pre-rRNA molecules had not been identified due to a lack of canonical RNA-binding domains. Recently, work by Piñeiro et al. showed AATF to be an RNAPI-associated RBP itself22. Their findings are in clear accordance with the fact that AATF has been identified in a row of RNA-interactome capture screens in different species^24,28,43^. A key finding of our study is the identification of the actual RNA-molecules bound by AATF using eCLIP with a clear overrepresentation of ribosomal RNAs. Furthermore, this approach allowed us to localize the binding sites in the 45S precursor molecule primarily to the cleavage sites required for SSU maturation, strengthening the hypothesized role of AATF in this process (Fig. 6). Several lines of published evidence also point towards this direction. Interestingly, Bfr2 – the budding yeast orthologue of AATF – was identified in a screen that analyzed constituents of yeast 90S particles assembled using truncated pre-18S RNAs and was shown to be specifically bound to the 5’ETS^40^. Furthermore, Bammert et al.^21^ had shown recently that loss of the ANN complex (containing AATF) resulted in 45S rRNA processing defects at the cleavage sites that are necessary for SSU generation. Our data provide strong indications that AATF is the actual RBP of this complex. However – just like AATF itself – both NGDN and NOL10 were detected in screens for polyA-tailed RNA-associated proteins and may possess the capacity to bind RNA themselves^28,29^. Indeed, Pineiro et al. showed NGDN binding to rRNA using RIP-qPCR^22^. As mentioned above, none of the ANN complex constituents – including AATF – possess a canonical RNA binding domain. While Bammert et al.^21^ argued that RNA binding of the ANN complex may be mediated by the WD40 domain of NOL10 which had previously been shown to mediate RNA-binding of other proteins^44^, the intrinsically disordered, basic C-terminus of AATF, 20% of which is made up of lysine (K) and arginine (R) residues, appears likely to promote RNA binding. This is also in line with recent studies that could emphasize the importance of intrinsically disordered regions (IDR) for RNA-protein interactions^24,27,45^ and the results of recent proteome-wide screens for RNA-binding domains^26,45^. Interestingly, work by He et al. showed that the two NoLS of AATF, the deletion of which resulted in a loss of rRNA binding in our hands, were directly associated with RNA^26^. Nevertheless, future experiments will be necessary to further dissect the exact molecular requirements of the physical interaction between AATF and RNA.

**Figure 6:**
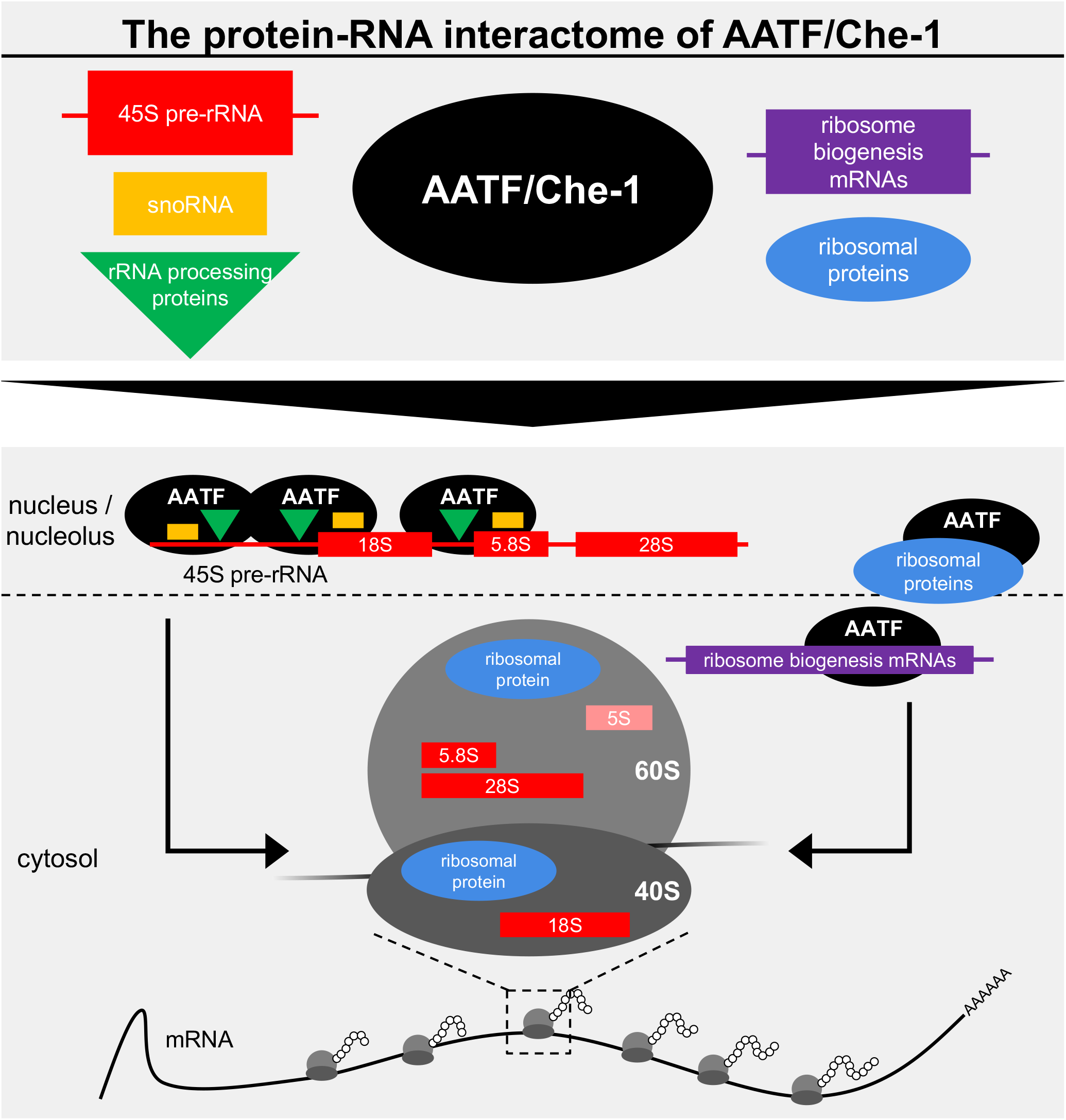
AATF – a key component of ribosome biogenesis. Using eCLIP and AP-MS, we establish that AATF interacts with several RNA species (45S pre-rRNA [red], snoRNAs [orange] and ribosome biogenesis mRNAs [purple]) and proteins (ribosomal [blue] and rRNA processing proteins [green]) involved in ribosome biogenesis. Through these interactions, AATF can recruit factors required for pre-rRNA processing to the actual cleavage sites. Additionally, AATF binds to ribosomal proteins and mRNAs encoding proteins known to play a role in ribosome assembly (right), which highlights the important role of AATF for ribosome biogenesis.

Our multilayer analysis - involving the global identification of the protein and RNA interactome - provides further evidence regarding the role of AATF in ribosome maturation. Beyond rRNA precursor molecules themselves AATF binds other RNA species important to the biogenesis of the ribosome. On the one hand, the mRNAs co-precipitating with AATF primarily encode proteins involved with RNA metabolism and ribosome maturation (Fig. 6). Here, it is intriguing to hypothesize that AATF may exert a regulatory function regarding the post-transcriptional regulation of these proteins. On the other hand, snoRNAs are highly enriched in the AATF interactome pointing towards a role of AATF in recruiting snoRNAs to pre-rRNA molecules (Fig. 6). The fact that AATF is associated with U3 snoRNA – considering that cleavage in the 5’ETS and the ITS1 of 45S pre-rRNA strongly depends on the U3 snoRNP – corroborates this hypothesis. This view is further complemented by the protein interactome containing numerous factors required for ribosome biogenesis. Since the vast majority of protein interactions – apart from the RBP SLIRP and the protein phosphatase PPP1CB – does not depend on RNA but appears to be mediated by direct protein-protein binding, AATF is likely to be involved in the coordination of protein-RNA supercomplexes in rRNA maturation (Fig. 6).

How is the molecular function of AATF in ribosome biogenesis linked to the previously described phenotypes regarding cellular proliferation and tumorigenesis? With rRNA availability being a central requirement for cellular survival and cell division, it is not surprising that loss of AATF in a knockout mouse model led to early embryonic lethality^31^. As to human disease this protein has been implicated to play an important role in cancer biology, a hypothesis that is partly based upon its ability to inhibit apoptosis. AATF has previously been found to be amplified or overexpressed in both hematological and solid tissue tumors, to correlate with poor prognosis, relapse and reduced survival^4–6,9,10,46^ and to mediate its effects on apoptosis by the modulation of p53 abundance and function^4,6^. Furthermore, increased ribosome biogenesis is not only employed by tumors to increase their proliferative potential but appears to be a risk factor for cancer onset on its own^47^. However, AATF appears to ensure cellular survival not only in the setting of cancer: its cytoprotective role has also been established in the setting of oxidative stress exposure to different cell types, including renal tubular cells and cardiomyocytes in models of acute kidney injury and ischemic reperfusion injury^48–50^. The known link between ribosome stress and p53 activity in the light of our new data allows for the exciting hypothesis that AATF mediates its effects on cell death employing this pathway. Based on the interaction between r-proteins and the E3 ubiquitin ligase MDM2, defects in ribosome maturation activate p53^19,36,51–53^. Loss of AATF would thus increase p53 activity linking the roles of AATF in ribosome biogenesis and p53 activity. In the light of an increasing number of studies trying to target ribosome biogenesis in this setting^54^ and taking into account that AATF has been shown to sensitize cancer but not normal cells to antineoplastic drugs^5^, future research shedding light on these aspects will be highly valuable.

## MATERIAL AND METHODS

### Molecular cloning and design of small interfering RNAs (siRNA)

For the generation of GFP-tagged AATF transgenes and transient overexpression of full-length AATF or 2ΔNoLS truncation, the AATF wild type sequence or the truncated version of it, generated by overlap extension PCR, were cloned into the modified AAV CAGGS GFP plasmid (Addgene #22212) or pcDNA6 (Invitrogen) using restriction enzymes. siRNA duplexes targeting the 3’ UTR of human AATF (accession number AJ249940.2) were custom-designed by Dharmacon.

### Cell culture, transfection and generation of single-copy transgenic cell lines using TALEN

Human HEK 293T and U2OS, and mouse IMCD-3 cells were grown in standard media at 37°C, 5% CO_2_ and routinely passaged using 0.05% Trypsin. Mycoplasma contamination was excluded using a commercial kit (Venor GeM, Sigma). Transfections for transient overexpression or stable integration of GFP-tagged transgenes were carried out on 60-80% confluent cells using calcium phosphate as described previously^55^, or lipofection (Lipofectamine 2000 and Lipofectamine LTX) and electroporation (Amaxa Nucleofector® kit V) according to the manufacturers’ instructions.

Transgenic cells were generated using TALEN plasmids targeting the AAVS1 locus (hAAVS1 1L TALEN, hAAVS1 1R TALEN and AAV-CAGGS-EGFP, all AddGene) as described previously^56,57^. 24h after transfection, cell lines were steadily selected with 2 μg/ml Puromycin. All cell lines were genotyped by integration PCR and phenotyped by both immunoblot and fluorescence light microscopy.

For knockdown experiments using commercial siRNA pools (Dharmacon), cell lines were transfected with Lipofectamine RNAiMAX and incubated for 48 h (final siRNA concentration 20 nM).

### eCLIP-seq, Read Processing and Cluster Analysis

eCLIP of AATF in K562 and HepG2 cells using primary anti-AATF antibody (Bethyl) was performed as previously described^58^. In brief, cells from two biological replicates were crosslinked using UV-C irradiation (254 nm, 150 mJ/cm^2^), subsequently lysed on ice and sonicated followed by DNA digestion and limited digestion of RNA using Turbo DNase (Thermo Scientific) and RNase I, respectively. A lysate aliquot was removed to serve as input control. Endogenous AATF was precipitated using primary anti-AATF antibody (Bethyl). Co-precipitated RNA was dephosphorylated and a 3’ RNA linker was ligated using T4 RNA ligase (NEB). The AATF-RNA complexes and input controls were run on a NuPage bis-tris protein gel, transferred to a nitrocellulose membrane and subsequently cut from the membrane to extract protein-bound RNA by Proteinase K digestion. RNA was cleaned using the RNA Clean & Concentrator kit (Zymo Research) and reverse transcribed using SuperScript III Reverse Transkrition kit (Thermo Scientific). After exonuclease digestion and alkaline phosphatase treatment (ExoSAP-IT, Affymetrix), a 3’ DNA adapter was ligated using the T4 RNA-ligase. The resulting cDNA library was amplified using Q5 Polymerase (NEB) and purified for sequencing analysis.

In addition to standard read processing and processing of reads to identify unique genomic mapping, reads mapping ribosomal RNA were quantified using a family-aware repeat element mapping pipeline that identifies reads unique to 45S pre-rRNA, 18S rRNA, or 28S rRNA respectively (Van Nostrand, E.L., *et al. in preparation*). To quantify relative enrichment between IP and input, relative information was calculated as the Kullback-Leibler divergence: 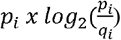, where *p_i_* is the fraction of total reads in IP that map to position *i*, and *q_i_* is the fraction of total reads in input for the same position. Regarding the definition of non-rRNA interaction partners we filtered the total dataset (K562 and HepG2 cells, 2 replicates each) for significant peaks over input (log_2_FC ≥3 and −log_10_ p-value ≥ 5) and collapsed all peaks mapping to one transcript to define a list of targets. For AATF eCLIP data accessibility refer to “Availability” section below.

To allow for a comparison of nucleolar to non-nucleolar RBPs, we assigned localization using manual curation of the RBP Image Database (http://rnabiology.ircm.qc.ca/RBPImage/). This resulted in 35 nucleolar proteins, 73 non-nucleolar proteins and 42 proteins that could not clearly be assigned (due to conflicting evidence in the literature) contained in the set of 150 RBPs of the ENCODE consortium^59^ (https://www.encodeproject.org/) that are the basis to the group of nucleolar proteins shown in the eCLIP analyses of this study (Suppl. Table 3).

### RIP-qPCR

HEK 293T cells were transfected with pcDNA6 plasmids containing sequences for triple FLAG-tagged wild-type AATF, a truncation lacking both nucleolar localization sites (AATF 2ΔNoLS), or red fluorescent protein (RFP). 24h after transfection the cells were washed twice with PBS and UV crosslinked (254 nm, 150 mJ/cm^2^) on ice. Following lysis in lysis buffer (50 mM Tris-HCl pH 7.4, 100 mM NaCl, 1% NP-40, 0.1% SDS, 0.5% sodium deoxycholate, Protease Inhibitor Cocktail III and Murine RNase Inhibitor) the samples were homogenized by passing three times through a 23G needle and sonicated (Bioruptor pico, 10x 30 sec ON/ 30 sec OFF) at 4°C. The samples were treated with Turbo-DNase (Thermo Scientific) at 37°C for 5 min and a lysate aliquot was removed to serve as input control. FLAG (M2) antibody was coupled to protein G dynabeads (Thermo Scientific). The RNA-protein complexes were immunoprecipitated over night at 4°C. Beads were washed three times with high salt buffer (50 mM Tris-HCl pH 7.4, 1 M NaCl, 1 mM EDTA, 1% NP-40, 0.1% SDS, 0.5% sodium deoxycholate) and five times with wash buffer (20 mM Tris-HCl pH 7.4, 10 mM MgCl_2_, 0.2% Tween-20). Subsequently, the RNA was recovered by TRIzol extraction and concentrated with RNA clean and concentrator columns (Zymo Research). Equal amounts of immunoprecipitated RNA were reverse transcribed using SuperScript III as described in the supplemental methods section and analysed by RT-qPCR. For quantitative analysis of 18S and 45S rRNA binding, the delta delta-CT method^60^ was applied to calculate the delta CT for full-length and mutant AATF as well as RFP (CT_IP_ − CT_INPUT_) and the subsequent delta delta-CT (e.g., delta CT_AATF_ − delta CT_RFP_) for three biological replicates.

### Co-immunoprecipitation and sample preparation for MS/MS

Seven 10 cm dishes (80% confluency) of HEK 293T Flp-In™ T-REx™ with stably integrated, inducible FLAG/HA-tagged AATF or GFP were used per replicate (n = 5). Cells were harvested in media, washed with ice-cold PBS and then lysed in modified RIPA buffer containing 1% NP-40 (IgPAL), 150 mM NaCl, 0.25% sodium-deoxycholate and 50 mM Tris and complete protease inhibitors (PIM; Roche). Lysates were sonicated (Bioruptor pico, 10x 30 sec ON/ 30 sec OFF), passed through a 23G syringe 3 times and subsequently cleared using centrifugation (16.000 g, 30 min at 4°C) and ultracentrifugation (210,000 g, 30 min at 4°C). The resulting supernatants were incubated with anti-FLAG beads (Miltenyi Biotec) for 2 h at 4°C. Co-immunoprecipitated proteins were isolated using magnetic μMACS columns (Miltenyi Biotec) as previously described^61^. For the investigation of RNA-dependency of the protein interactions, RNA was digested on-column by adding 1 U RNase I and 25 U Benzonase (in wash buffer) for 30 min at 4°C. After three washing steps, samples were eluted from the column in Urea 2M buffer and subsequently reduced (5 mM DTT, 30 min, 55°C) and alkylated (40 mM CAA, 30 min, 20°C in the dark). Proteins were digested at 37°C for 16 h using trypsin and lysC (both at a 1:75 enzyme-protein ratio) following standard protocols. Formic acid was added to a final concentration of 1% to stop proteolysis. Peptides were loaded onto StageTips, desalted and labeled on column using triplex dimethyl labelling^62^. Labels were shuffled between replicates to exclude any bias. After elution, light, medium and heavy-labelled peptides from one experiment were pooled, dried down in a vacuum concentrator. Peptides were stored in 5% DMSO / 1% formic acid at −20°C prior LC-MS analysis ^63,64^. For details on MS processing see Suppl. Methods.

### MS data analysis (protein interactome)

All mass spectrometric raw data were processed with Maxquant (version 1.5.3.8) using default parameters. Protein quantification was performed based on MS1-level peptide ion intensities. Maxquant first determines peptide ion intensities, which are used to calculate peptide ratios between light/medium/heavy labeled peptides. Protein ratios are derived from the mean of all associated peptide ratios and subsequently used for statistical testing. This is described in detail in the Maxquant publication^65^. Briefly, MS2 spectra were searched against the Uniprot HUMAN.fasta database, including a list of common contaminants. False discovery rates on protein and PSM level were estimated by the target-decoy approach to 1% (Protein FDR) and 1% (PSM FDR) respectively. The minimal peptide length was set to 7 amino acids and carbamidomethylation at cysteine residues was considered as a fixed modification. Oxidation (M) and Acetyl (Protein N-term) were included as variable modifications. The match-between runs option was disabled. Dimethyl triplex labeling quantification was used, and the re-quantify option was enabled. Maxquant output files were further processed using Perseus (version 1.5.5.3) and R/Bioconductor. Obtained protein ratios were log_2_ transformed. Proteins flagged as “only identified by site”, “reverse” and “potential contaminant” were removed from the data set. In total, 5 biological replicates were analyzed (no technical replicates). Labels were shuffled between replicates to exclude any bias. Statistical significance of putative AATF interactors was assessed utilizing a one-sided t-test (fudge factor S0 = 1)^66^, using the log_2_ transformed ratios of AATF vs GFP pulldown (Fig. 4A/D, Suppl. Fig.4A/D). For comparison of protein abundance in AATF pulldowns with and without RNase the log_2_ ratios were normalized such that the ratio of AATF in those two conditions equals zero, assuming that AATF itself remains unaffected by the RNase treatment. For this comparison, a two-sided t-test was used (S0 = 0.1) to determine statistical significance. For both data sets, q-values were determined using the method of Benjamini and Hochberg.

### Endogenous co-immunoprecipitation

For immunoprecipitation of endogenous AATF, one 10 cm dish of HEK 293T cells were used. Cells were harvested in PBS, resuspended in modified RIPA buffer (50 mM Tris pH 7.5, 150 mM NaCl, 1% Igepal, 0,25% Na-Deoxycholate, protease inhibitor cocktail without EDTA-Roche), homogenized through a 23G needle on ice, sonicated (BioRuptor Pico, 10x 30 sec ON, 30 sec OFF) and incubated with 2 U/ml Turbo DNase (Invitrogen) for 10 min at 37°C, shaking. Immunoprecipitation with anti-AATF antibody (2 μg, Sigma) was performed as described above, with rabbit IgG serving as negative control, using magnetic Dynabeads Potein G beads (ThermoFisher Scientific). After overnight incubation, beads were concentrated on-magnet, washed five times with modified RIPA buffer and boiled with equal amounts of SDS loading buffer. Samples were analyzed by Western blotting (Suppl. methods).

## DATA AVAILABILITY

AATF K562 eCLIP data has been deposited at the ENCODE Data Coordination Center (https://www.encodeproject.org/) under accession identifier ENCSR819XBT, and HepG2 data has been deposited at the Gene Expression Omnibus (series GSE107766; samples GSM2878484 (Rep 1), GSM2878485 (Rep 2), GSM2878554 (size-matched input)). The Interactome dataset has been uploaded in ProteomeXchange via the PRIDE database (https://www.ebi.ac.uk/pride/archive/). Project accession: PXD011055; These data will be made publicly accessible upon publication.

## ACCESSION NUMBERS

Protein interactome PRIDE database: Project accession: PXD011055 eCLIP data GEO: series GSE107766; samples GSM2878484, GSM2878485, GSM2878554 eCLIP data ENCODE: ENCSR819XBT

## ACKNOWLEDGEMENTS

We thank Stefanie Keller, Serena Greco-Torres, Martyna Brütting and Ruth Herzog for excellent technical assistance. Special thanks to Constantin Rill for help with TALEN transgenesis. The monoclonal antibody E7 developed by M. McCutcheon and S. Carroll was obtained from the Developmental Studies Hybridoma Bank, created by the NICHD of the NIH and maintained at The University of Iowa, Department of Biology, Iowa City, IA 52242.

## AUTHOR CONTRIBUTIONS

F.F. and R.-U.M. designed the study; R.K., M.I., E.L.V.N., M.J., S.C., H.H., M.S., K.H., P.K., L.S., and F.F. performed experiments; R.K., M.I., E.L.V.N., C.F., K.Bu., M. Ho., K.Bo., P.F., R.-U.M., G.W.Y., C.D., M.He. and F.F. analyzed the data; R.K., M.I., F.F. and R.-U.M. prepared the figures; R.-U.M., R.K., M.I., F.F., B.S. and T.B. drafted and revised the paper; all authors approved the final version of the manuscript.

## REFERENCES

1 Bruno, T. et al. Che-1 affects cell growth by interfering with the recruitment of HDAC1 by Rb. Cancer Cell 2, 387–399 (2002).

2 Fanciulli, M. et al. Identification of a novel partner of RNA polymerase II subunit 11, Che-1, which interacts with and affects the growth suppression function of Rb. FASEB J 14, 904–912 (2000).

3 Ishigaki, S. et al. AATF mediates an antiapoptotic effect of the unfolded protein response through transcriptional regulation of AKT1. Cell Death Differ 17, 774–786, doi:10.1038/cdd.2009.175 (2010).

4 Welcker, D. et al. AATF suppresses apoptosis, promotes proliferation and is critical for Kras-driven lung cancer. Oncogene 37, 1503–1518, doi:10.1038/s41388-017-0054-6 (2018).

5 Höpker, K. et al. AATF/Che-1 acts as a phosphorylation-dependent molecular modulator to repress p53-driven apoptosis. EMBO J 31, 3961–3975, doi:10.1038/emboj.2012.236 (2012).

6 Bruno, T. et al. Che-1 promotes tumor cell survival by sustaining mutant p53 transcription and inhibiting DNA damage response activation. Cancer Cell 18, 122–134, doi:10.1016/j.ccr.2010.05.027 (2010).

7 Bruno, T. et al. Che-1 phosphorylation by ATM/ATR and Chk2 kinases activates p53 transcription and the G2/M checkpoint. Cancer Cell 10, 473–486, doi:10.1016/j.ccr.2006.10.012 (2006).

8 Desantis, A. et al. Che-1 modulates the decision between cell cycle arrest and apoptosis by its binding to p53. Cell Death Dis 6, e1764 doi:10.1038/cddis.2015.117 (2015).

9 Bruno, T. et al. Che-1 sustains hypoxic response of colorectal cancer cells by affecting Hif-1α stabilization. J Exp Clin Cancer Res 36, 32, doi:10.1186/s13046-017-0497-1 (2017).

10 Desantis, A. et al. Che-1-induced inhibition of mTOR pathway enables stress-induced autophagy. EMBO J 34, 1214–1230, doi:10.15252/embj.201489920 (2015).

11 Bruno, T., Iezzi, S. & Fanciulli, M. Che-1/AATF: A Critical Cofactor for Both Wild-Type- and Mutant-p53 Proteins. Front Oncol 6, 34, doi:10.3389/fonc.2016.00034 (2016).

12 Iezzi, S. & Fanciulli, M. Discovering Che-1/AATF: a new attractive target for cancer therapy. Front Genet 6, 141, doi:10.3389/fgene.2015.00141 (2015).

13 Ferraris, S. E. et al. Nucleolar AATF regulates c-Jun-mediated apoptosis. Mol Biol Cell 23, 4323–4332, doi:10.1091/mbc.E12-05-0419 (2012).

14 Henras, A. K., Plisson-Chastang, C., O’Donohue, M. F., Chakraborty, A. & Gleizes, P. E. An overview of pre-ribosomal RNA processing in eukaryotes. Wiley Interdiscip Rev RNA 6, 225–242, doi:10.1002/wrna.1269 (2015).

15 Tafforeau, L. et al. The complexity of human ribosome biogenesis revealed by systematic nucleolar screening of Pre-rRNA processing factors. Mol Cell 51, 539–551, doi:10.1016/j.molcel.2013.08.011 (2013).

16 Badertscher, L. et al. Genome-wide RNAi Screening Identifies Protein Modules Required for 40S Subunit Synthesis in Human Cells. Cell Rep 13, 2879–2891, doi:10.1016/j.celrep.2015.11.061 (2015).

17 Nicolas, E. et al. Involvement of human ribosomal proteins in nucleolar structure and p53-dependent nucleolar stress. Nat Commun 7, 11390, doi:10.1038/ncomms11390 (2016).

18 Steitz, T. A. A structural understanding of the dynamic ribosome machine. Nat Rev Mol Cell Biol 9, 242–253, doi:10.1038/nrm2352 (2008).

19 Liu, Y., Deisenroth, C. & Zhang, Y. RP-MDM2-p53 Pathway: Linking Ribosomal Biogenesis and Tumor Surveillance. Trends Cancer 2, 191–204, doi:10.1016/j.trecan.2016.03.002 (2016).

20 Boulon, S., Westman, B. J., Hutten, S., Boisvert, F. M. & Lamond, A. I. The nucleolus under stress. Mol Cell 40, 216–227, doi:10.1016/j.molcel.2010.09.024 (2010).

21 Bammert, L., Jonas, S., Ungricht, R. & Kutay, U. Human AATF/Che-1 forms a nucleolar protein complex with NGDN and NOL10 required for 40S ribosomal subunit synthesis. Nucleic Acids Res, doi:10.1093/nar/gkw790 (2016).

22 Pineiro, D. et al. Identification of the RNA polymerase I-RNA interactome. Nucleic Acids Res, doi:10.1093/nar/gky779 (2018).

23 Ignarski, M. et al. The RNA-Protein Interactome of Differentiated Kidney Tubular Epithelial Cells. J Am Soc Nephrol, doi:10.1681/ASN.2018090914 (2019).

24 Beckmann, B. M. et al. The RNA-binding proteomes from yeast to man harbour conserved enigmRBPs. Nat Commun 6, 10127, 10.1038/ncomms10127 [doi] (2015).

25 Liao, Y. et al. The Cardiomyocyte RNA-Binding Proteome: Links to Intermediary Metabolism and Heart Disease. Cell Rep 16, 1456–1469, 10.1016/j.celrep.2016.06.084 [doi] (2016).

26 He, C. et al. High-Resolution Mapping of RNA-Binding Regions in the Nuclear Proteome of Embryonic Stem Cells. Mol Cell 64, 416–430, doi:10.1016/j.molcel.2016.09.034 (2016).

27 Hentze, M. W., Castello, A., Schwarzl, T. & Preiss, T. A brave new world of RNA-binding proteins. Nat Rev Mol Cell Biol, doi:10.1038/nrm.2017.130 (2018).

28 Castello, A. et al. Insights into RNA biology from an atlas of mammalian mRNA-binding proteins. Cell 149, 1393–1406, doi:10.1016/j.cell.2012.04.031 (2012).

29 Baltz, A. G. et al. The mRNA-bound proteome and its global occupancy profile on protein-coding transcripts. Mol Cell 46, 674–690, doi:10.1016/j.molcel.2012.05.021 (2012).

30 Consortium, E. P. A user’s guide to the encyclopedia of DNA elements (ENCODE). PLoS Biol 9, e1001046, doi:10.1371/journal.pbio.1001046 (2011).

31 Thomas, T., Voss, A. K., Petrou, P. & Gruss, P. The murine gene, Traube, is essential for the growth of preimplantation embryos. Dev Biol 227, 324–342, doi:10.1006/dbio.2000.9915 (2000).

32 Scott, M. S., Troshin, P. V. & Barton, G. J. NoD: a Nucleolar localization sequence detector for eukaryotic and viral proteins. BMC Bioinformatics 12, 317, doi:10.1186/1471-2105-12-317 (2011).

33 Lafontaine, D. L. & Tollervey, D. The function and synthesis of ribosomes. Nat Rev Mol Cell Biol 2, 514–520, doi:10.1038/35080045 (2001).

34 Phipps, K. R., Charette, J. & Baserga, S. J. The small subunit processome in ribosome biogenesis-progress and prospects. Wiley Interdiscip Rev RNA 2, 1–21, doi:10.1002/wrna.57 (2011).

35 Scott, M. S. & Ono, M. From snoRNA to miRNA: Dual function regulatory non-coding RNAs. Biochimie 93, 1987–1992, doi:10.1016/j.biochi.2011.05.026 (2011).

36 Lindstrom, M. S. et al. Nucleolus as an emerging hub in maintenance of genome stability and cancer pathogenesis. Oncogene, doi:10.1038/s41388-017-0121-z (2018).

37 Yoshihama, M., Nakao, A. & Kenmochi, N. snOPY: a small nucleolar RNA orthological gene database. BMC Res Notes 6, 426, doi:10.1186/1756-0500-6-426 (2013).

38 Yoshimoto, R., Kataoka, N., Okawa, K. & Ohno, M. Isolation and characterization of post-splicing lariat-intron complexes. Nucleic Acids Res 37, 891–902, doi:10.1093/nar/gkn1002 (2009).

39 Brannan, K. W. et al. SONAR Discovers RNA-Binding Proteins from Analysis of Large-Scale Protein-Protein Interactomes. Mol Cell 64, 282–293, 10.1016/j.molcel.2016.09.003 [doi] (2016).

40 Soltanieh, S., Lapensee, M. & Dragon, F. Nucleolar proteins Bfr2 and Enp2 interact with DEAD-box RNA helicase Dbp4 in two different complexes. Nucleic Acids Res 42, 3194–3206, doi:10.1093/nar/gkt1293 (2014).

41 Barandun, J., Hunziker, M. & Klinge, S. Assembly and structure of the SSU processome-a nucleolar precursor of the small ribosomal subunit. Curr Opin Struct Biol 49, 85–93, doi:10.1016/j.sbi.2018.01.008 (2018).

42 Calo, E. et al. RNA helicase DDX21 coordinates transcription and ribosomal RNA processing. Nature 518, 249–253, doi:10.1038/nature13923 (2015).

43 Baltz, A. G. et al. The mRNA-bound proteome and its global occupancy profile on protein-coding transcripts. Mol Cell 46, 674–690, doi:10.1016/j.molcel.2012.05.021 (2012).

44 Loedige, I. et al. The Crystal Structure of the NHL Domain in Complex with RNA Reveals the Molecular Basis of Drosophila Brain-Tumor-Mediated Gene Regulation. Cell Rep 13, 1206–1220, doi:10.1016/j.celrep.2015.09.068 (2015).

45 Castello, A. et al. Comprehensive Identification of RNA-Binding Domains in Human Cells. Mol Cell 63, 696–710, doi:10.1016/j.molcel.2016.06.029 (2016).

46 Folgiero, V. et al. Che-1 is targeted by c-Myc to sustain proliferation in pre-B-cell acute lymphoblastic leukemia. EMBO Rep 19, doi:10.15252/embr.201744871 (2018).

47 Derenzini, M., Montanaro, L. & Trere, D. Ribosome biogenesis and cancer. Acta Histochem 119, 190–197, doi:10.1016/j.acthis.2017.01.009 (2017).

48 Xie, J. & Guo, Q. Apoptosis antagonizing transcription factor protects renal tubule cells against oxidative damage and apoptosis induced by ischemia-reperfusion. J Am Soc Nephrol 17, 3336–3346, doi:10.1681/ASN.2006040311 (2006).

49 Wang, D., Chen, T. Y. & Liu, F. J. Che-1 attenuates hypoxia/reoxygenation-induced cardiomyocyte apoptosis by upregulation of Nrf2 signaling. Eur Rev Med Pharmacol Sci 22, 1084–1093, doi:10.26355/eurrev_201802_14395 (2018).

50 Guo, S. et al. Che-1 inhibits oxygen-glucose deprivation/reoxygenation-induced neuronal apoptosis associated with inhibition of the p53-mediated proapoptotic signaling pathway. Neuroreport, doi:10.1097/WNR.0000000000001095 (2018).

51 Donati, G. et al. Selective inhibition of rRNA transcription downregulates E2F-1: a new p53-independent mechanism linking cell growth to cell proliferation. J Cell Sci 124, 3017–3028, doi:10.1242/jcs.086074 (2011).

52 Ofir-Rosenfeld, Y., Boggs, K., Michael, D., Kastan, M. B. & Oren, M. Mdm2 regulates p53 mRNA translation through inhibitory interactions with ribosomal protein L26. Mol Cell 32, 180–189, doi:10.1016/j.molcel.2008.08.031 (2008).

53 Ogawa, L. M. & Baserga, S. J. Crosstalk between the nucleolus and the DNA damage response. Mol Biosyst 13, 443–455, doi:10.1039/c6mb00740f (2017).

54 Drygin, D. et al. Targeting RNA polymerase I with an oral small molecule CX-5461 inhibits ribosomal RNA synthesis and solid tumor growth. Cancer Res 71, 1418–1430, doi:10.1158/0008-5472.CAN-10-1728 (2011).

55 Chen, C. & Okayama, H. High-efficiency transformation of mammalian cells by plasmid DNA. Mol Cell Biol 7, 2745–2752 (1987).

56 Hockemeyer, D. et al. Efficient targeting of expressed and silent genes in human ESCs and iPSCs using zinc-finger nucleases. Nat Biotechnol 27, 851–857, doi:10.1038/nbt.1562 (2009).

57 Sanjana, N. E. et al. A transcription activator-like effector toolbox for genome engineering. Nat Protoc 7, 171–192, doi:10.1038/nprot.2011.431 (2012).

58 Van Nostrand, E. L. et al. Robust transcriptome-wide discovery of RNA-binding protein binding sites with enhanced CLIP (eCLIP). Nat Methods 13, 508–514, 10.1038/nmeth.3810 [doi] (2016).

59 Eric L Van Nostrand, P. F., Gabriel A Pratt, Xiaofeng Wang, Xintao Wei, Rui Xiao, Steven M Blue, Jia-Yu Chen, Neal A.L. Cody, Daniel Dominguez, Sara Olson, Balaji Sundararaman, Lijun Zhan, Cassandra Bazile, Louis Philip Benoit Bouvrette, Julie Bergalet, Michael O Duff, Keri E. Garcia, Chelsea Gelboin-Burkhart, Myles Hochman, Nicole J Lambert, Hairi Li, Thai B Nguyen, Tsultrim Palden, Ines Rabano, Shashank Sathe, Rebecca Stanton, Amanda Su, Ruth Wang, Brian A. Yee, Bing Zhou, Ashley L Louie, Stefan Aigner, Xiang-dong Fu, Eric Lécuyer, Christopher B. Burge, Brenton R. Graveley, Gene W. Yeo. A Large-Scale Binding and Functional Map of Human RNA Binding Proteins. bioRxiv, doi:https://doi.org/10.1101/179648 (2018).

60 Livak, K. J. & Schmittgen, T. D. Analysis of relative gene expression data using real-time quantitative PCR and the 2(-Delta Delta C(T)) Method. Methods 25, 402–408, doi:10.1006/meth.2001.1262 (2001).

61 Hubner, N. C. & Mann, M. Extracting gene function from protein-protein interactions using Quantitative BAC InteraCtomics (QUBIC). Methods 53, 453–459, doi:10.1016/j.ymeth.2010.12.016 (2011).

62 Boersema, P. J., Raijmakers, R., Lemeer, S., Mohammed, S. & Heck, A. J. Multiplex peptide stable isotope dimethyl labeling for quantitative proteomics. Nat Protoc 4, 484–494, doi:10.1038/nprot.2009.21 (2009).

63 Kohli, P. et al. Label-free quantitative proteomic analysis of the YAP/TAZ interactome. American journal of physiology. Cell physiology 306, C805–818, doi:10.1152/ajpcell.00339.2013 (2014).

64 Rappsilber, J., Ishihama, Y. & Mann, M. Stop and go extraction tips for matrix-assisted laser desorption/ionization, nanoelectrospray, and LC/MS sample pretreatment in proteomics. Analytical chemistry 75, 663–670 (2003).

65 Cox, J. & Mann, M. MaxQuant enables high peptide identification rates, individualized p.p.b.-range mass accuracies and proteome-wide protein quantification. Nat Biotechnol 26, 1367–1372, doi:10.1038/nbt.1511 (2008).

66 Tusher, V. G., Tibshirani, R. & Chu, G. Significance analysis of microarrays applied to the ionizing radiation response. Proc Natl Acad Sci U S A 98, 5116–5121, doi:10.1073/pnas.091062498 (2001).

67 Huang da, W., Sherman, B. T. & Lempicki, R. A. Systematic and integrative analysis of large gene lists using DAVID bioinformatics resources. Nat Protoc 4, 44–57, doi:10.1038/nprot.2008.211 (2009).

