## Supplemental Material and Methods for "A protein-RNA interaction atlas of the ribosome biogenesis factor AATF"

**Supplemental Tables**……………………………………………separate .xls spreadsheets

**Supplemental Tables and Figures**

**Supplemental Figure 1.** (related to figure 1)
**AATF is a conserved RNA-binding protein localizing to the nucleolus and interacting with rRNA**

**Supplemental Figure 2.** (related to figure 2)
**Comparison of the rRNA-binding sites of AATF to other nucleolar and non-nucleolar RBPs in ENCODE**

**Supplemental Figure 3.** (related to figure 3)
**Impact of AATF depletion on rRNA modifications**

**Supplemental Figure 4.** (related to figure 4)
**RNA binding proteins, R-proteins and RNAPI RNA interactors are part of the AATF protein interactome**

**Supplemental Figure 5.**
**Original blots and images**

**Supplemental Table 1.**
**The AATF RNA interactome (after rRNA removal) as identified by eCLIP**

**Supplemental Table 2.**
**The AATF protein interactome as identified by MS/MS**

**Supplemental Table 3.** (related to figure 1A and suppl. figure 2)
**List of all RBPs contained in the ENCODE datasets and their (non-)nucleolar designation**

### Supplemental material and methods

#### Primary Antibodies

| targeted protein | manufacturer | order ID |
|---|---|---|
| AATF | Abnova | H00026574-A01 |
| AATF | Sigma-Aldrich | HPA004940 |
| AATF | Bethyl | A301-032A lot 001 |
| FLAG (M2) | Sigma-Aldrich | F1804-1MG |
| GFP (B2) | SantaCruz BioTech | sc-9996 |
| HEATR1 | SantaCruz BioTech | sc-390445 |
| FBL | GeneTex | GTX24566 |
| NOP2 (NSUN1) | SantaCruz BioTech | sc-398884 |
| TUBB/β-Tubulin (E7) | Develop. Studies Hybridoma Bank | E7 |

#### DNA/RNA oligonucleotides

| Genotyping | Sequence 5´-3´ |
|---|---|
| AAVS1 Integration PCR HA-R 2R | FP GTGAGTTTGCCAAGCAGTCA |
| AAVS1 Integration PCR HA-R 2F | RP TATCCGCTCACAATTCCACA |
| AAVS 1 Empty locus | RP CGGAACTCTGCCCTCTAACG |

| RT-qPCR | |
|---|---|
| **Target** | **Sequence 5´-3´** |
| 18S* | FP CTCAACACGGGAAACCTCAC<br>RP CGCTCCACCAACTAAGAACG |
| AATF | FP CTTGGACACGGACAAAAGGT<br>RP CTCCAGACCCTTCCTCATCA |
| ACTB | FP GGACTTCGAGACAAGAGATGG<br>RP AGCACTGTGTTGGCGTACAG |
| HPRT | FP TGACACTGGCAAAACAATGCA<br>RP GGTCCTTTTCACCAGCAAGCT |
| 45S | FP ACCCACCCTCGGTGAGA<br>RP CAAGGCACGCCTCTCAGAT |

* While the primers amplifying 45S pre-rRNA were designed to exclusively amplify this precursor transcript, amplification of the precursors by the 18S rRNA primers cannot be excluded, since 18S rRNA does not contain internal splice or other processing sites that would enable transcript-specific discrimination.

| RNAi | | | |
|---|---|---|---|
| ON-TARGETplus Human AATF siRNA SMARTpool | CAAGCGCUCUGUCUAUCGA, GUGAUGACCUUCUCUAGUG, GCACUUAAAGCAUUGUUGA, CCAGGGUGAUUGACAGGUU | Dharmacon | L-004373-00-0005 |
| ON-TARGETplus Mouse AATF siRNA SMARTpool | GACACGAGACAUAUAGUAA, GUGAGUAGCAUUAGAAAGU, CUAUAGGAAUCACACACUA, GGACGGAGUUGUUUCGAUC | Dharmacon | L-050461-00-0005 |
| ON-TARGETplus Non-targeting pool | GUUUACAUGUCGACUAA, UGGUUUACAUGUUGUGUGA, UGGUUUACAUGUUUUCUGA, UGGUUUACAUGUUUUCCUA | Dharmacon | D-001810-10-05 |

| | | | |
|---|---|---|---|
| Human AATF Custom siRNA (3' UTR specific) | GAGCAUUGUUACCGCCAAAUU, UUUGGCGGUAACAAUGCUCUU | Dharmacon | - |

**Western blot**

Cells were washed once with ice-cold PBS, lysed on ice in either iCLIP lysis buffer or modified RIPA buffer for 15 minutes and then sonicated (BioRuptor Pico, 10x 30 sec ON, 30 sec OFF). Lysates were then run on PAGE gels, transferred onto PVDF membranes and blocked in 5% bovine serum albumin. Primary antibodies were incubated either for 1h at room temperature or overnight at 4°C. Secondary HRP coupled antibodies (Jackson ImmunoResearch, 1:20,000 dilution) were incubated for 1h at room temperature. Bands were visualized using ECL reagents, at a Fusion chemoluminescence imaging system (Peqlab) (see table above for a list of primary antibodies used).

**Fluorescence light microscopy**

Cells were grown on coverslips coated with 100 µg/ml Collagen I to 60-70% confluency. Cells were then washed once with PBS supplemented with $Ca^{2+}/Mg^{2+}$ (PBS+), fixed with 4% PFA for 5 minutes and rinsed three times with PBS+ and mounted on ProLong Gold mounting medium with DAPI (Invitrogen) to visualize nuclei. Images were acquired using a Zeiss epifluorescence microscope (Zeiss Axiovert 200M).

**RNA isolation and RT-qPCR**

Whole RNA was isolated from all cells using TRIzol® and subsequent chloroform extraction or a commercial column-based kit (ZymoResearch) according to the manufacturers' instructions. RNA for RNAseq experiments or modification analysis was DNase digested on-column. Both oligo(dT) primers and random hexamers were used for reverse transcription with either SuperScript III Reverse Transcription kit or High Capacity cDNA Reverse Transcription kit (both Thermo Scientific). Real-time quantitative PCR was performed using either SYBR Green Master Mix (Applied Biosystems) or commercial probe-based assays (Thermo Scientific, IDT) on either an Applied Biosystems 7900HT Fast Real-Time PCR System or an Applied Biosystems QuantStudio™ 12K Flex Real-Time PCR System. All experiments were performed on biological and technical triplicates. For the analysis of miRNA expression upon knockdown of AATF, TaqMan® MicroRNA Arrays A and B (Applied Biosystems) were used according to the manufacturer's instructions. Cycle thresholds were normalized to housekeeping genes such as ACTB or GAPDH using the delta-delta-CT method as described previously [2]. For all primer sequences, refer to Suppl. Methods.

**Statistical analysis**

Unless stated otherwise, a two-tailed, unpaired student's t-test was used for statistical analysis of e.g. immunofluorescence data using GraphPad Prism v5. Statistical significance is indicated by asterisks as follows: *p<0,05, **p<0,01, ***p<0.,05, ****p<0,001, ns = not significant. Depicted results show the mean of at least three independent experiments/biological replicates. Unless stated otherwise, error bars depict the standard deviation of the mean.

**GO Term analysis**

Gene ontology (GO) analyses were performed using the Database for Annotation, Visualization and Integrated Discovery (DAVID) v6.8 (https://david.ncifcrf.gov/)[3]. Online results were downloaded as .txt- files and depicted using GraphPad Prism v5 or Microsoft Excel.

**LC-MS analysis (protein interactome)**

All samples were analyzed on a Q Exactive Plus (Thermo Scientific) that was coupled to an EASY nLC 1200 (Thermo Scientific). Peptides were loaded with solvent A (0.1% formic acid in water) onto an in-house packed analytical column (50 cm, 75 µm I.D., filled with 2.7 µm Poroshell EC120 C18, Agilent). Peptides were chromatographically separated at a constant flow rate of 250 nL/min using the following gradient: 10-23% solvent B (0.1% formic acid in 80 % acetonitrile) within 75.0 min, 23-39% solvent B within 5.0 min, 39-95% solvent B within 5.0 min, followed by washing and column equilibration. The mass spectrometer was operated in data-dependent acquisition mode. The MS1 survey scan was acquired from 300-1750 m/z at a resolution of 70,000. The top 10 most abundant peptides were isolated within a 1.8 Th window and subjected to HCD fragmentation at a normalized collision energy of 27%. The AGC target was set to 5e5 charges, allowing a maximum injection time of 120 ms. Product ions were detected in the Orbitrap at a resolution of 35,000. Precursors were dynamically excluded for 20.0s.

**Analysis of RNA modifications: LC-MS/MS**

**RNA hydrolysis to nucleoside level**

The RNA samples were digested into nucleosides as it was described before[6]: 500 ng of RNA were treated with 0.3 U nuclease P1 (Roche Diagnostics, Germany), 0.1 U snake venom phosphodiesterase (Worthington, USA), 200 ng Pentostatin (Adenosine deaminase inhibitor, Sigma-Aldrich, Germany) and 500 ng Tetrahydrouridine (Cytidine deaminase inhibitor, Merck-Millipore, Germany) in 1/10 vol. of 10x nuclease P1 buffer (10x NP1 Buffer: 9 vol 250 mM $NH_4OAc$, pH 5.0; 1 vol 2 mM $ZnCl_2$) for 2 h at 37 °C. Next, 1/10 vol. of 10x fast alkaline phosphatase buffer (10x FAP Buffer: 100 mM M $NH_4OAc$, pH 9.0) and 1 U fast alkaline phosphatase (Fermentas, Germany) were added and the mixture was incubated at 37 °C for another 60 min.

For LC-MS/MS analysis 3x 25 ng of each RNA sample were employed (technical triplicates).

**Relative quantification of modified nucleosides *via* LC-MS/MS**

For RNA analysis, an Agilent1260 series equipped with a diode array detector (DAD) and a Triple Quadrupole mass spectrometer (Agilent 6460) were utilized. In addition to that, a Synergy Fusion RP18 column (4 µm particle size, 80 Å pore size, 250 mm length, 2 mm inner diameter) from Phenomenex (Germany) was used at 35°C and separation of nucleosides was performed using a flow rate of 0.35 mL/min. 5 mM ammonium acetate buffer (pH 5.3) served as solvent A and acetonitrile (LCMS grade, Sigma-Aldrich, Germany) as solvent B. The respective LC-gradient is shown in table M1 below.

**Table M1:** Gradient for nucleoside separation prior to MS analysis.

| Time [min] | Solvent A [%] | Solvent B[%] |
|---|---|---|
| 0 | 100 | 0 |
| 10 | 92 | 8 |

| | | |
|---|---|---|
| 20 | 60 | 40 |
| 23 | 100 | 0 |
| 30 | 100 | 0 |

The main nucleosides (cytidine, uridine, guanosine, adenosine) were measured photometrically at 254 nm by the DAD and modified nucleosides were analyzed *via* the mass spectrometer (operated in the positive ion mode) equipped with an electrospray ion source (Agilent Jet Stream, for settings see table M2).

**Table M2:** Electrospray ionization settings.

| **Parameter** | |
|---|---|
| Gas temperature | 350 °C |
| Gas flow | 8 L/min |
| Nebulizer pressure | 50 psi |
| Sheath gas temperature | 350 °C |
| Sheath gas flow | 12 L/min |
| Capillary voltage | 3000 V |

To monitor the mass transitions of the modified nucleosides, Agilent Mass Hunter software was used in the Dynamic Multiple Reaction Monitoring (DMRM) mode. Further details are displayed in Table M3.

**Table M3:** MS-parameters of the Dynamic Multiple Reaction Monitoring mode.

| **Modified nucleoside** | **Molecular weight [Da]** | **Precursor ion [m/z]** | **Product ion [m/z]** | **Retention time [min]** | **Fragmentor voltage [V]** | **Collision energy [eV]** | **Cell accelerator voltage [V]** |
|---|---|---|---|---|---|---|---|
| $A_m$ | 281 | 282 | 136 | 16 | 92 | 13 | 2 |
| $^{13}C/^{15}N$-$A_m$ | 292 | 293 | 141 | 16 | 92 | 13 | 2 |
| $C_m$ | 257 | 258 | 112 | 9.5 | 60 | 9 | 2 |
| $^{13}C/^{15}N$-$C_m$ | 270 | 271 | 119 | 9.5 | 60 | 9 | 2 |
| $G_m$ | 297 | 298 | 152 | 12.6 | 72 | 5 | 2 |
| $^{13}C/^{15}N$-$G_m$ | 313 | 314 | 162 | 12.6 | 72 | 5 | 2 |
| $m^1A$ | 281 | 282 | 150 | 5.6 | 92 | 17 | 2 |
| $^{13}C/^{15}N$-$m^1A$ | 297 | 298 | 161 | 5.6 | 92 | 17 | 2 |
| $m^2_2G$ | 311 | 312 | 180 | 14.9 | 82 | 9 | 2 |
| $^{13}C/^{15}N$-$m^2_2G$ | 328 | 329 | 192 | 14.9 | 82 | 9 | 2 |
| $m^6_2A$ | 295 | 296 | 164 | 18.6 | 102 | 17 | 2 |
| $^{13}C/^{15}N$-$m^6_2A$ | 312 | 313 | 176 | 18.6 | 102 | 17 | 2 |
| $m^5C$ | 257 | 258 | 126 | 8.8 | 40 | 9 | 2 |
| $^{13}C/^{15}N$-$m^5C$ | 270 | 271 | 134 | 8.8 | 40 | 9 | 2 |
| $m^6A$ | 281 | 282 | 150 | 16.7 | 92 | 17 | 2 |
| $^{13}C/^{15}N$-$m^6A$ | 297 | 298 | 161 | 16.7 | 92 | 17 | 2 |
| $m^7G$ | 297 | 298 | 166 | 8.7 | 82 | 9 | 2 |
| $^{13}C/^{15}N$-$m^7G$ | 313 | 314 | 177 | 8.7 | 82 | 9 | 2 |

| | | | | | | | |
|---|---|---|---|---|---|---|---|
| Ψ | 244 | 245 | 209 | 4 | 81 | 5 | 2 |
| $^{13}C/^{15}N$- Ψ | 255 | 256 | 220 | 4 | 81 | 5 | 2 |
| $U_m$ | 258 | 259 | 113 | 11.1 | 66 | 5 | 2 |
| $^{13}C/^{15}N$- $U_m$ | 270 | 271 | 119 | 11.1 | 66 | 5 | 2 |

Resulting spectra were processed using Agilent MassHunter Qualitative Analysis Software: In a first step, the recorded UV chromatogram of the main nucleoside guanosine was extracted to receive the 'area under the curve' (AUC) and then, the UV-signal derived from the SIL-IS (see below) was subtracted. After that, calibration measurements of guanosine dilutions (5-500 pmol) were applied for exact quantification and the amount of injected guanosine (in pmol) of each RNA sample was calculated by using the resulting guanosine calibration factor.

Quantification of modified nucleosides was achieved by utilizing $^{13}C$- and $^{15}N$-labeled total RNA from *C. elegans* as a Stable Isotope-Labeled Internal Standard (SIL-IS)[7]: For each investigated modification, 10 calibration solutions (0.1-5000 fmol) consisting of the modified reference nucleoside (all Sigma-Aldrich, Germany) including 20 ng SIL-IS were prepared and subjected to LC-MS measurement. The resulting mass spectra of each solution were then processed by integrating the MS/MS peaks to receive the AUC values. In the next step, the ratios of the extracted peak areas of the modified nucleosides and their corresponding $^{13}C$-labeled isotopes were calculated and plotted using Microsoft Excel to receive calibration curves. This enabled the determination of the slopes of these curves which corresponds to the nucleoside-isotope response factors for each modification. To obtain the modification content in the RNA samples, the same amount of SIL-IS (= 20 ng) was added and again the resulting ratio of modified nucleoside and its $^{13}C$-labeled isotope was calculated. Afterwards, the latter was divided by the corresponding modification response factor to receive the modification amount in fmol. Lastly, the ratios of modified nucleosides and the injected guanosine amounts were calculated. Further and more detailed information can be found in Thüring et al.[7] and Kellner et al.[6].

**Table M4: List of modifications**

| **Abbreviation** | **Name** |
|---|---|
| $A_m$ | 2′-O-methyladenosine |
| $C_m$ | 2′-O-methylcytidine |
| $G_m$ | 2′-O-methylguanosine |
| $m^1A$ | 1-methyladenosine |
| $m^2_2G$ | *N2,N2*-dimethylguanosine |
| $m^6_2A$ | *N6,N6*-dimethyladenosine |
| $m^5C$ | 5-methylcytidine |
| $m^6A$ | *N6*-methyladenosine |
| $m^7G$ | 7-methylguanosine |
| Ψ | pseudouridine |
| $U_m$ | 2′-O-methyluridine |

### Supplemental Figure legends

**Suppl. Figure 1: AATF is a conserved RNA-binding protein localizing to the nucleolus and interacting with rRNA**

**A** Graphical representation of AATF enrichment comparing our dataset (striped)[8] to published mRNA interactome datasets (black)[9-14]. Both human AATF and its highly conserved mouse and yeast orthologues (Aatf/Traube and Brefeldin A-resistance protein 2/Bfr2, respectively) show a significant enrichment across data sets. The $Log_2$ fold change was calculated for crosslinked over non-crosslinked samples.

**B** Stepwise scheme of the eCLIP-protocol. 1. Intact cells were UV crosslinked and lysed. 2. Representation of crosslink formed between protein and RNA after exposure to UV light. 3. The cell lysate was submitted to partial RNase I digestion. 4. The RBP-RNA complex was immunoprecipitated. 5. On the beads, the RNA was dephosphorylated and a 3´ RNA adapter was ligated. 6. The lysate was size selected via PAGE and the RNA was extracted. 7. Reverse transcription was performed with the reaction stopping at the site of crosslinking after which the RNA was removed. 8. A 3´DNA adapter was ligated and the fragments were PCR amplified. 9. The fragments were Illumina indexed and submitted to paired-end sequencing.

**C** The amino acid sequence of AATF was analyzed for nucleolar localization sequences (NoLS) using the Nucleolar localization sequence Detector tool (NoD)[15], where the AATF sequence (UNIPROT ID Q9NY61) was plotted against a NoLS likelihood score. Two NoLSs (as assessed by a score of > 0.8) were identified in the C-terminal part of the protein: AA 326-345 (NoLS 1) and AA 494-522 (NoLS 2).

**D** Scheme of protein domains and sequence features of wild type AATF (LZ: leucin zipper, NoLS: nucleolar localization signal) and a truncated version of the protein lacking both nucleolar localization signal.

**E** Localisation of WT and mutant AATF. Fluorescence images of transgenic U2OS cell lines expressing either stably integrated full-length GFP-AATF (upper panels) or the GFP-tagged AATF 2ΔNoLS truncation (lower panels). Deletion of both nucleolar localization sequences results in displacement of the protein from the nucleolus. Nuclei were counterstained with DAPI. DIC: differential interference contrast.

**F** Western blot related to RIP-qPCR analysis of 18S rRNA and 45S pre-rRNA (Fig. 1D). Western blot analysis (anti-FLAG antibody) shows efficient precipitation of FLAG-tagged proteins, which were expressed and precipitated at the same level. FLAG-RFP (red fluorescent protein) served as negative control.

**G** U2OS cells were transfected with either an siRNA pool targeting the AATF coding sequence or non-targeted control siRNA. After 48h, cells were harvested and RNA was isolated. qPCR for ACTB, AATF and 18S rRNA on RNA derived from these cells confirms the significant decrease of 18S rRNA after AATF depletion.

**Suppl. Figure 2: Comparison of the RNA-binding sites of AATF to other nucleolar and non-nucleolar RBPs in ENCODE**

Heatmaps indicate the relative information (IP versus input) for ribosomal RNA18S, RNA28S, and RNA45S transcripts. Groupings indicate 2 replicates for AATF, 35 nucleolar RBPs, and 73 non-nucleolar datasets. Nucleolar and non-nucleolar localization of RBPs was based on ENCODE I data (see methods section).

**Suppl. Figure 3: Impact of AATF depletion on rRNA modifications**

Analysis of rRNA modifications of whole RNA isolated from murine IMCD cells after transfection with AATF-targeted siRNA or a control using LC-MS/MS (see suppl. methods for details). Here, no differences between control cells and cells depleted of AATF were detected for the most common modifications.

**Suppl. Figure 4: RNA binding proteins, R-proteins and RNAPI RNA interactors are part of the AATF protein interactome**

**A** Scatter plot showing AATF protein interactors annotated as RNA binding proteins. According to the comparison with known RBPs about 80% of the proteins interacting with AATF are themselves annotated as RNA-binding proteins (red dots: known RBPs, black dot: AATF, white dots: AATF interacting proteins not annotated as RBPs).
**B** Bar chart depicting R-proteins that are part of the (partially) non-RNA- and RNA-dependent protein interactome of AATF. The grey lower part of the bars depict the percentage of known R-proteins[16] in the analyzed subgroups.
**C** Venn diagram depicting the high level of overlap of the AATF protein interactome as well as the RNAPI RNA interactome as published by Piñeiro et al.[17].
**D** Scatter plot showing that some of the AATF interactors were shown in the recent publication by Pinero et al. to be RNA binding proteins either dependent or independent.on RNAPI inhibition Dark green: RNAPI independent interactors, light green: RNAPI dependent interactors, white dots: not identified in RNAPI RNA interactome.

**Suppl. Figure 5: Original blots and images**

### Supplemental Table legends

**Suppl. Table 1: The AATF RNA interactome (after rRNA removal) as identified by eCLIP**

**A: HepG2, replicate 1, annotated .bed files**

**B: HepG2, replicate 2, annotated .bed files**

**C: K562, replicate 1, annotated .bed files**

**D: K562, replicate 2, annotated .bed files**

Each dataset was filtered for significant peaks FC>=3 and p-value>=5, and pooled.

To define a list of bound RNAs, replicates ENSG identifiers were removed

The ENSG IDs were then annotated for Gene Stable ID, Gene name and Gene Type using biomart on the ENSEMBL website (ensembl genes 93) 17 IDs were not annotated (obsolete)

**E: "bound RNAs"**, 665 targets with at least one significant peak in 1 experiment (FC≥3 and p-value≥5) in 1 out of the 4 experiments (2 replicates K562 cells, 2 replicates HepG2 cells)

**F: "at least in 2 experiments",** 292 targets all sequences containing a significant peak (FC>=3 and p-value>=5) in at least 2 out of the 4 experiments (2 replicates K562 cells, 2 replicates HepG2 cells)

**Suppl. Table 2: The AATF protein interactome as identified by MS/MS**

**"data":** all proteins identified in all biological replicates (n = 5).

**"A-AATF bona fide interactors":** only proteins that show $\log_2$FC >=2 and $-\log_{10}$ pval >=1.3 in AATF vs. GFP

**"B-RNA dependent":** all proteins among bona fide interactors that show a $\log_2$FC <=-2 and $-\log_{10}$ pval >=1.3 in AATF+RNAse vs AATF normalized

**"C-partially RNA dependent":** all proteins among bona fide interactors that show a $\log_2$FC <=-1 and $-\log_{10}$ pval >=1.0 in AATF+RNAse vs AATF normalized

**"D-partially non-RNA dependent":** all proteins among bona fide interactors that show a $\log_2$FC >=1 and $-\log_{10}$ pval >=1.0 in AATF RNAse vs. GFP

**"E-non RNA dependent":** all proteins among bona fide interactors that show a $\log_2$FC >=2.0 and $-\log_{10}$ pval >=1.3 in AATF RNAse vs. GFP

**"F-AATF bona fide vs Pineiro":** all proteins in A matched versus the RNApol I RNA interactome (3)

**Supplemental Table 3.** (related to figure 1A and suppl. figure 2): **ENCODE RBPs list**

**"A_total ENCODE list":** list of 150 proteins from the ENCODE consortium for which eCLIP datasets are available - 223 datasets in total, in 2 different cell lines (HepG2 and K562).

**"B_nucleolar"** and **"C_non nucleolar":** subsets of RBPs were chosen according to their localization based mostly on immunofluorescence information from the website: http://rnabiology.ircm.qc.ca/RBPImage/.

All proteins for which insufficient or contradictory data were available to support either localization or biological function, were removed from this analysis.

### References

1 Schindelin, J. *et al.* Fiji: an open-source platform for biological-image analysis. *Nat Methods* **9**, 676-682, doi:10.1038/nmeth.2019 (2012).

2 Livak, K. J. & Schmittgen, T. D. Analysis of relative gene expression data using real-time quantitative PCR and the 2(-Delta Delta C(T)) Method. *Methods* **25**, 402-408, doi:10.1006/meth.2001.1262 (2001).

3 Huang da, W., Sherman, B. T. & Lempicki, R. A. Systematic and integrative analysis of large gene lists using DAVID bioinformatics resources. *Nat Protoc* **4**, 44-57, doi:10.1038/nprot.2008.211 (2009).

4 Cox, J. & Mann, M. MaxQuant enables high peptide identification rates, individualized p.p.b.-range mass accuracies and proteome-wide protein quantification. *Nat Biotechnol* **26**, 1367-1372, doi:10.1038/nbt.1511 (2008).

5 Tusher, V. G., Tibshirani, R. & Chu, G. Significance analysis of microarrays applied to the ionizing radiation response. *Proc Natl Acad Sci U S A* **98**, 5116-5121, doi:10.1073/pnas.091062498 (2001).

6 Kellner, S. *et al.* Absolute and relative quantification of RNA modifications via biosynthetic isotopomers. *Nucleic Acids Res* **42**, e142, doi:10.1093/nar/gku733 (2014).

7 Thüring, K., Schmid, K., Keller, P. & Helm, M. Analysis of RNA modifications by liquid chromatography-tandem mass spectrometry. *Methods* **107**, 48-56, doi:10.1016/j.ymeth.2016.03.019 (2016).

8 Ignarski, M. *et al.* The RNA-Protein Interactome of Differentiated Kidney Tubular Epithelial Cells. *J Am Soc Nephrol*, doi:10.1681/ASN.2018090914 (2019).

9 Liao, Y. *et al.* The Cardiomyocyte RNA-Binding Proteome: Links to Intermediary Metabolism and Heart Disease. *Cell Rep* **16**, 1456-1469, doi:S2211-1247(16)30853-1 [pii]10.1016/j.celrep.2016.06.084 [doi] (2016).

10 He, C. *et al.* High-Resolution Mapping of RNA-Binding Regions in the Nuclear Proteome of Embryonic Stem Cells. *Mol Cell* **64**, 416-430, doi:10.1016/j.molcel.2016.09.034 (2016).

11 Beckmann, B. M. *et al.* The RNA-binding proteomes from yeast to man harbour conserved enigmRBPs. *Nat Commun* **6**, 10127, doi:ncomms10127 [pii]10.1038/ncomms10127 [doi] (2015).

12 Hentze, M. W., Castello, A., Schwarzl, T. & Preiss, T. A brave new world of RNA-binding proteins. *Nat Rev Mol Cell Biol*, doi:10.1038/nrm.2017.130 (2018).

13 Castello, A. *et al.* Insights into RNA biology from an atlas of mammalian mRNA-binding proteins. *Cell* **149**, 1393-1406, doi:10.1016/j.cell.2012.04.031 (2012).

14 Baltz, A. G. *et al.* The mRNA-bound proteome and its global occupancy profile on protein-coding transcripts. *Mol Cell* **46**, 674-690, doi:10.1016/j.molcel.2012.05.021 (2012).

15 Scott, M. S., Troshin, P. V. & Barton, G. J. NoD: a Nucleolar localization sequence detector for eukaryotic and viral proteins. *BMC Bioinformatics* **12**, 317, doi:10.1186/1471-2105-12-317 (2011).

16 Nicolas, E. *et al.* Involvement of human ribosomal proteins in nucleolar structure and p53-dependent nucleolar stress. *Nat Commun* **7**, 11390, doi:10.1038/ncomms11390 (2016).

17 Pineiro, D. *et al.* Identification of the RNA polymerase I-RNA interactome. *Nucleic Acids Res*, doi:10.1093/nar/gky779 (2018).

**Suppl. Figure 1** (relates to Fig.1)

A

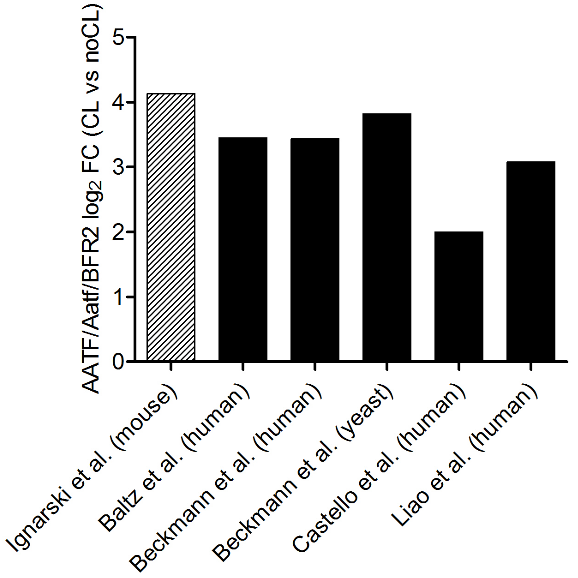

B

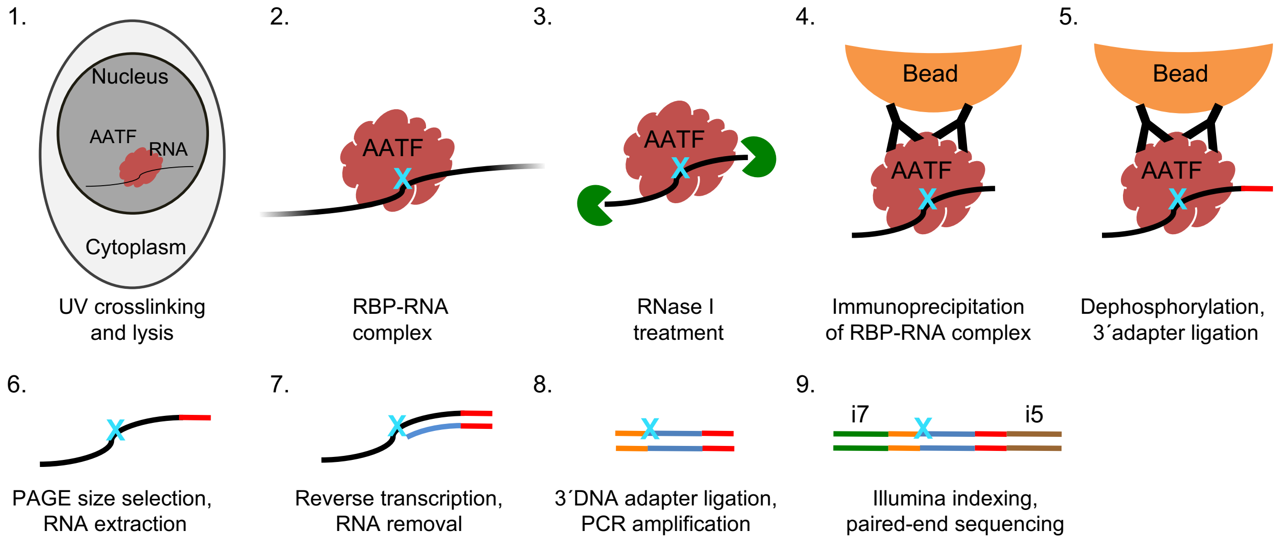

C

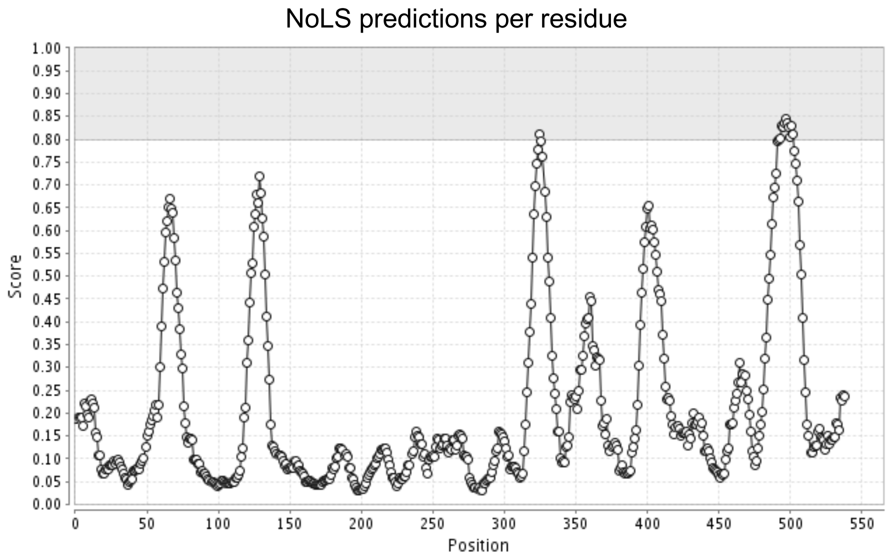

**Suppl. Figure 1** (relates to Fig.1)

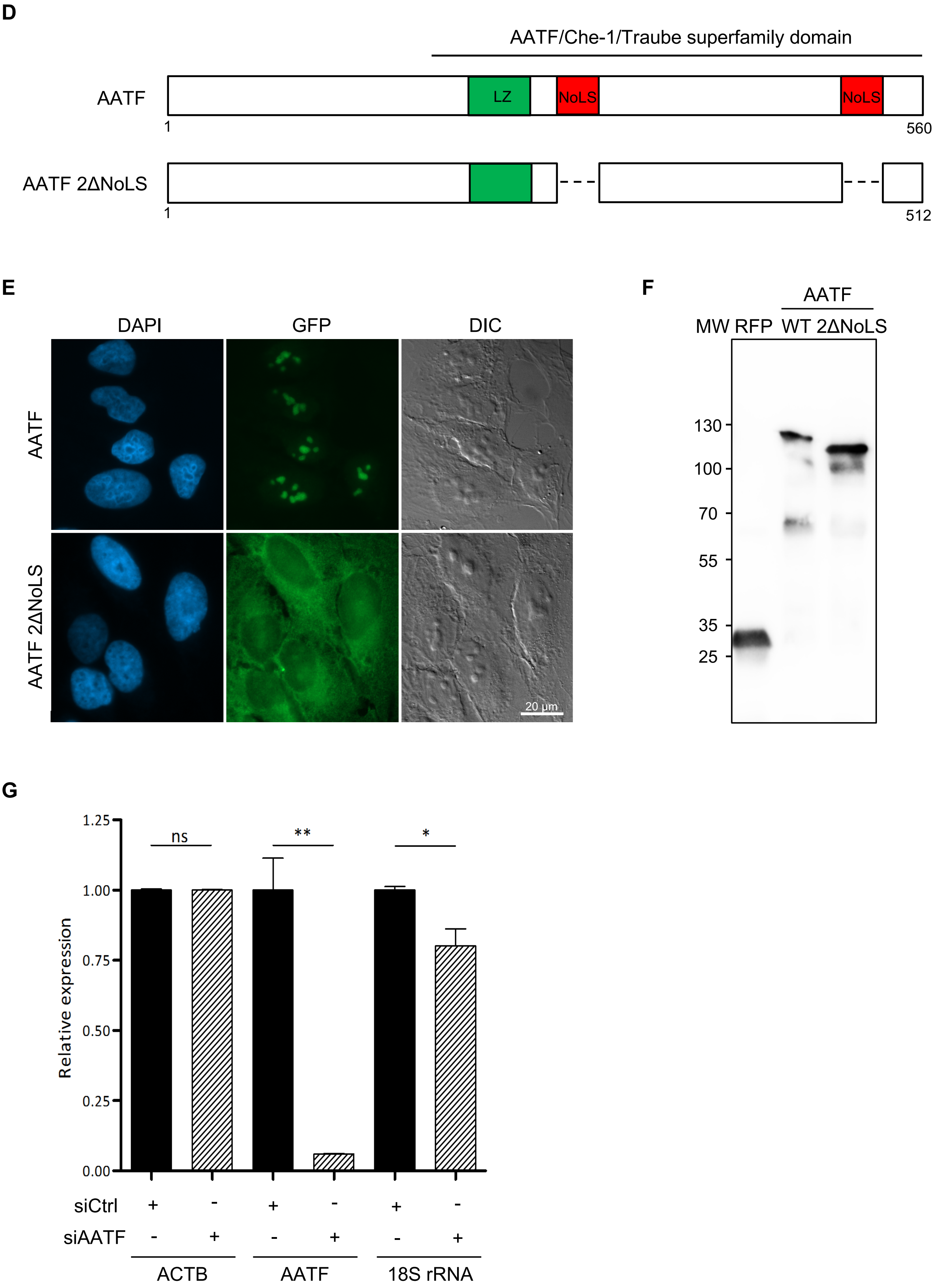

**Suppl. Figure 2** (relates to Fig. 2)

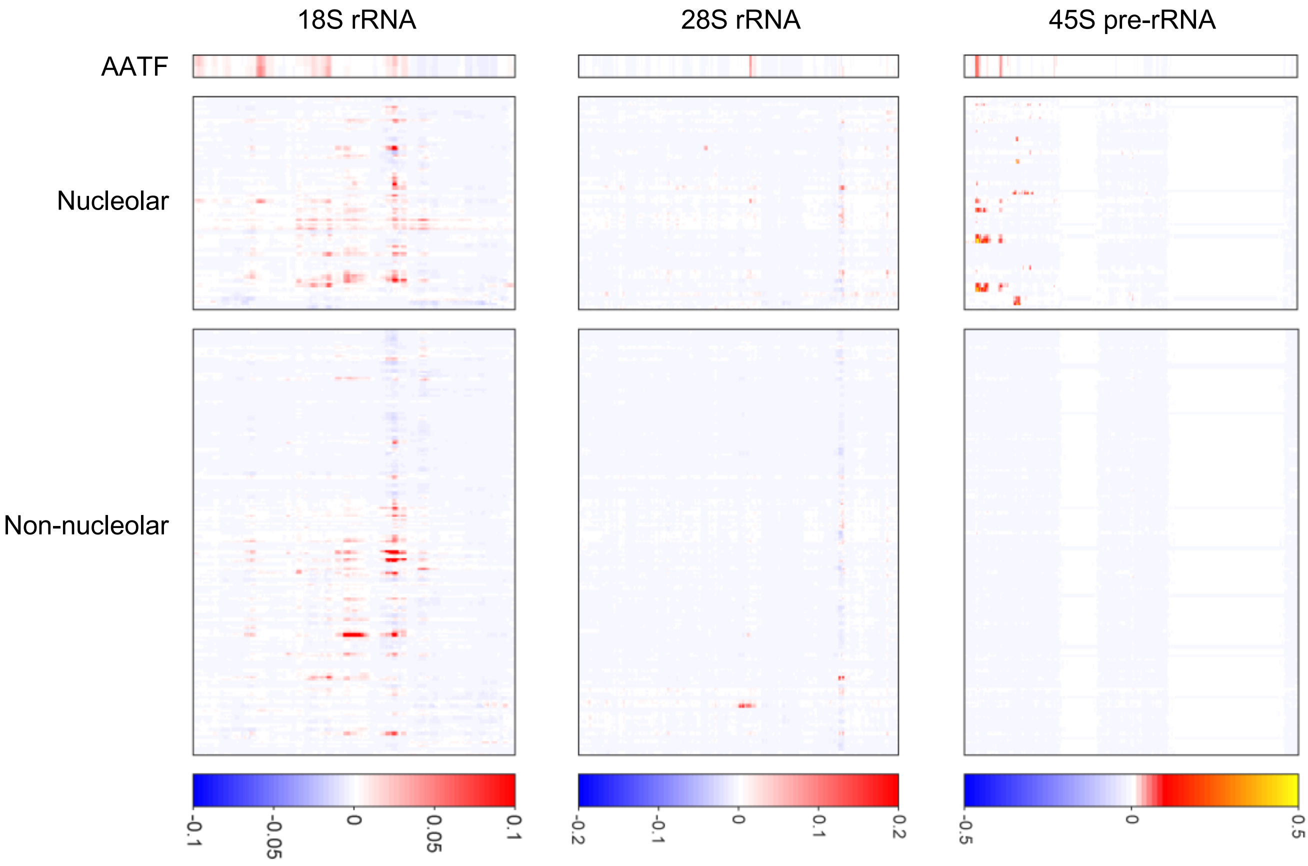

**Suppl. Figure 3** (relates to Fig. 3)

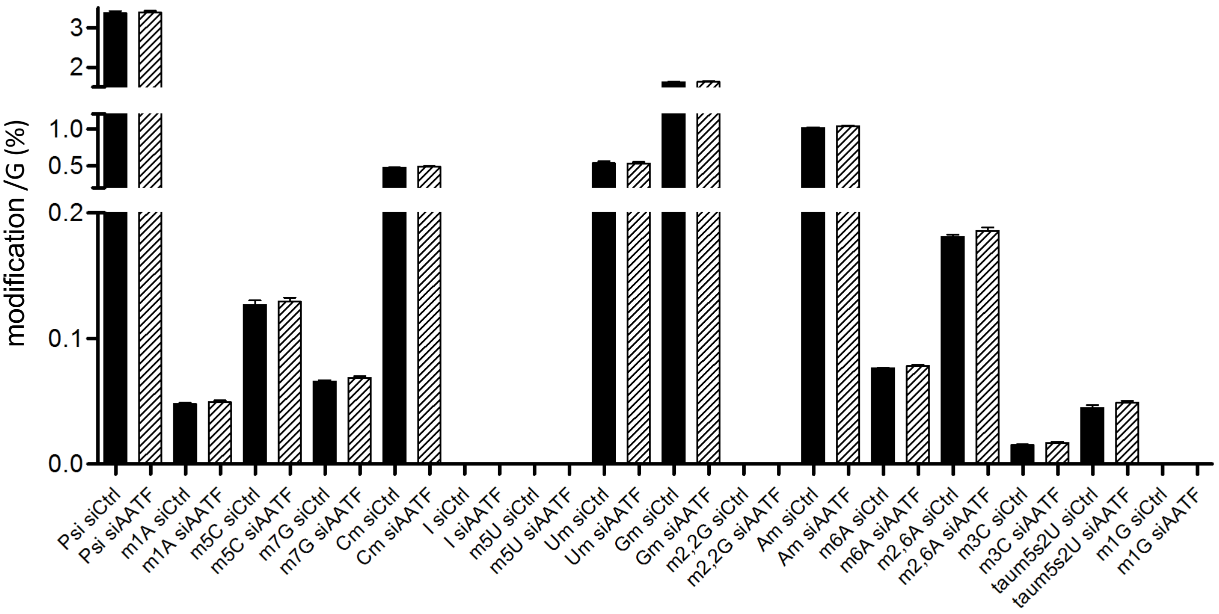

**Suppl. Figure 4** (relates to Fig. 4)

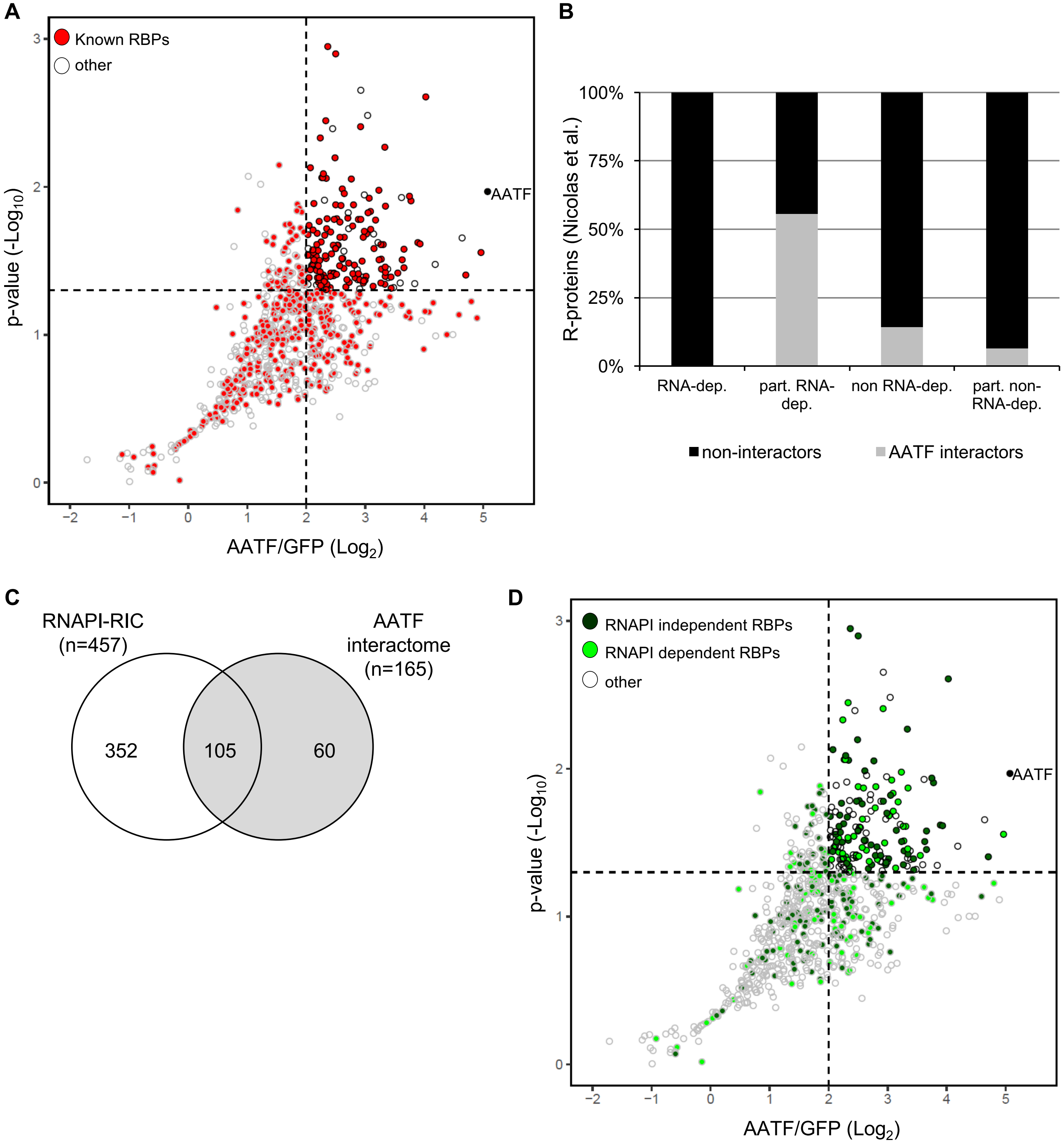

**Suppl. Figure 5** (original blots and images)

**A** (relates to Fig. 1D)

**anti-AATF**

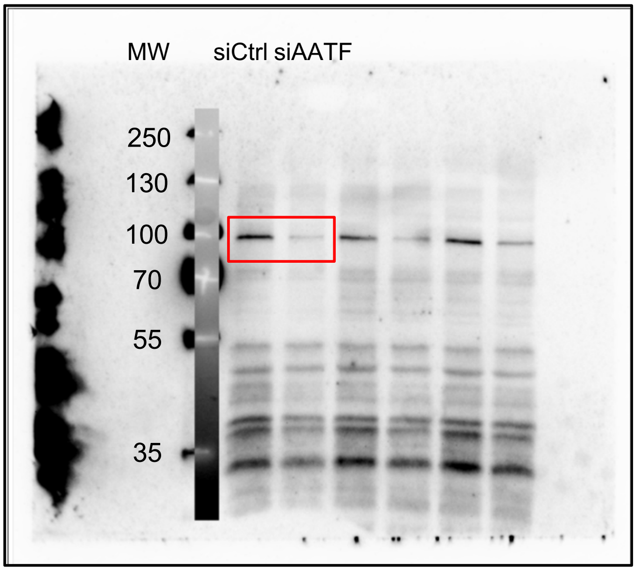

**anti-β-tubulin**

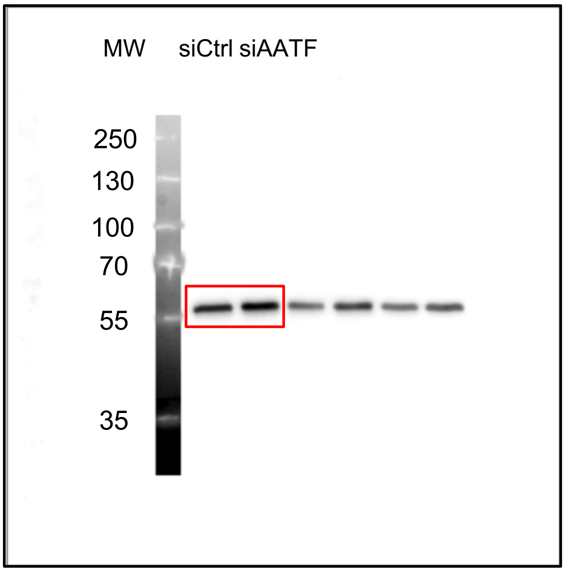

**EtBr stained agarose gel**

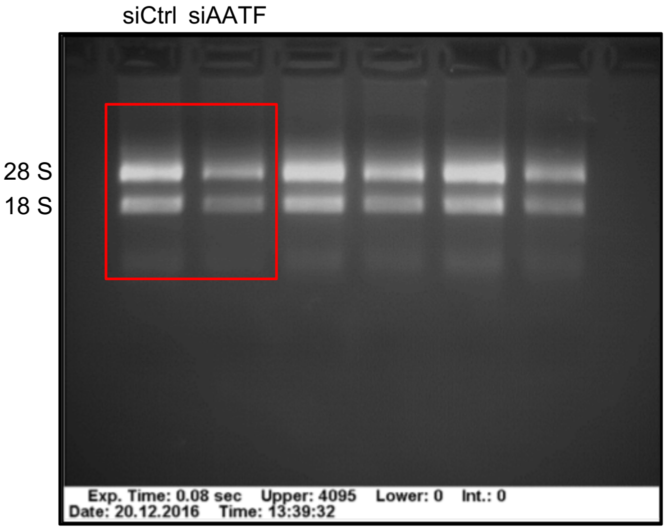

**B** (relates to Fig. 1E)

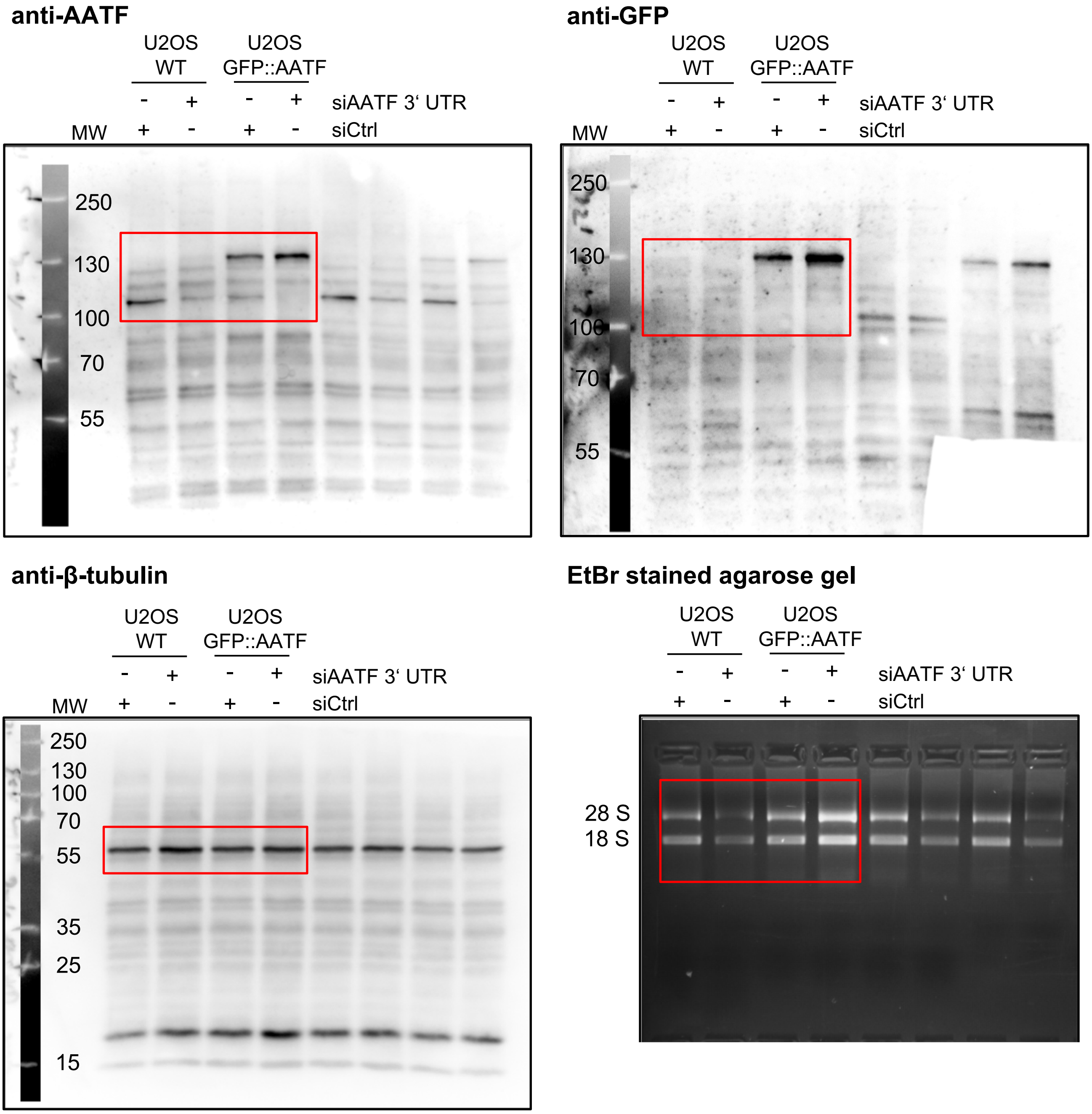

**C** (relates to suppl. Fig. 1E)

**AATF**

DAPI GFP

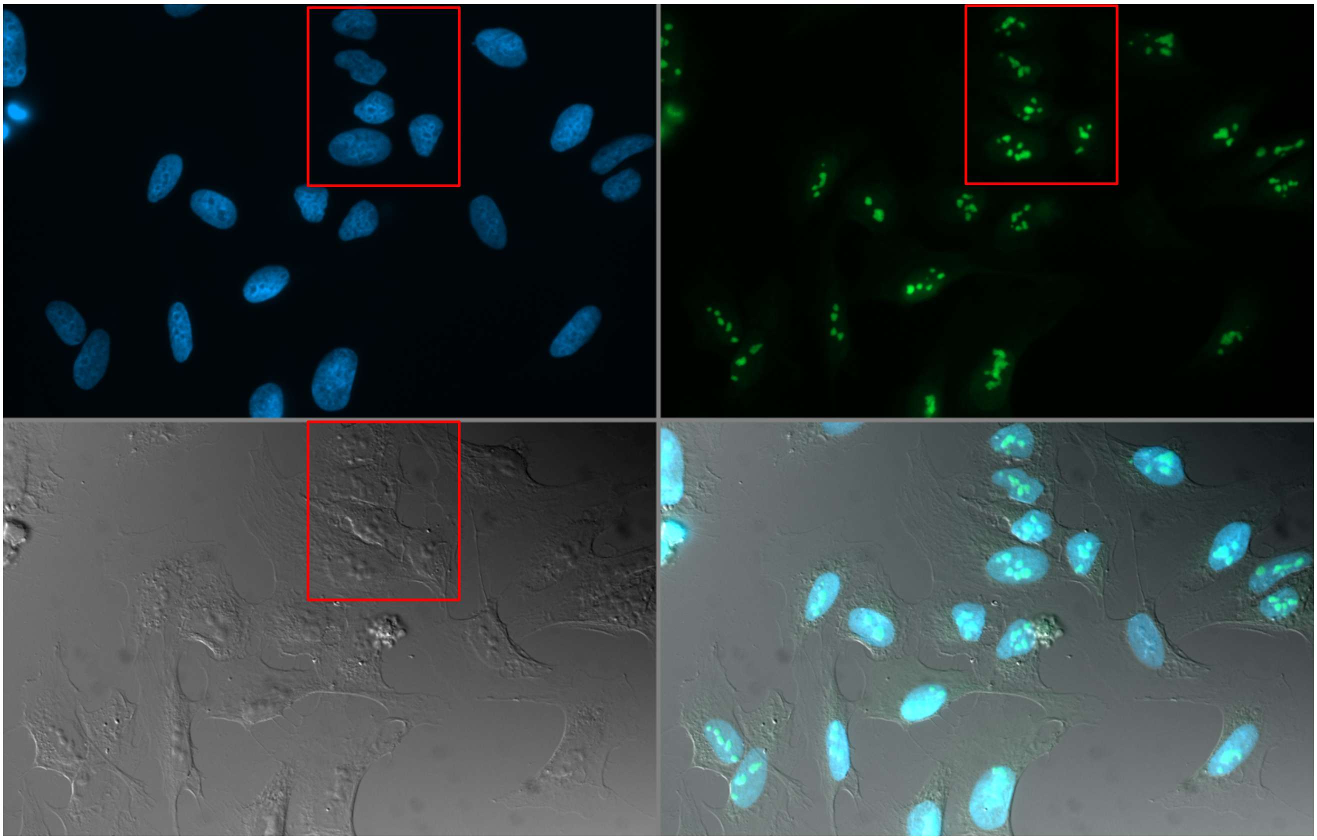

DIC Merge

**AATF 2ΔNoLS**

DAPI GFP

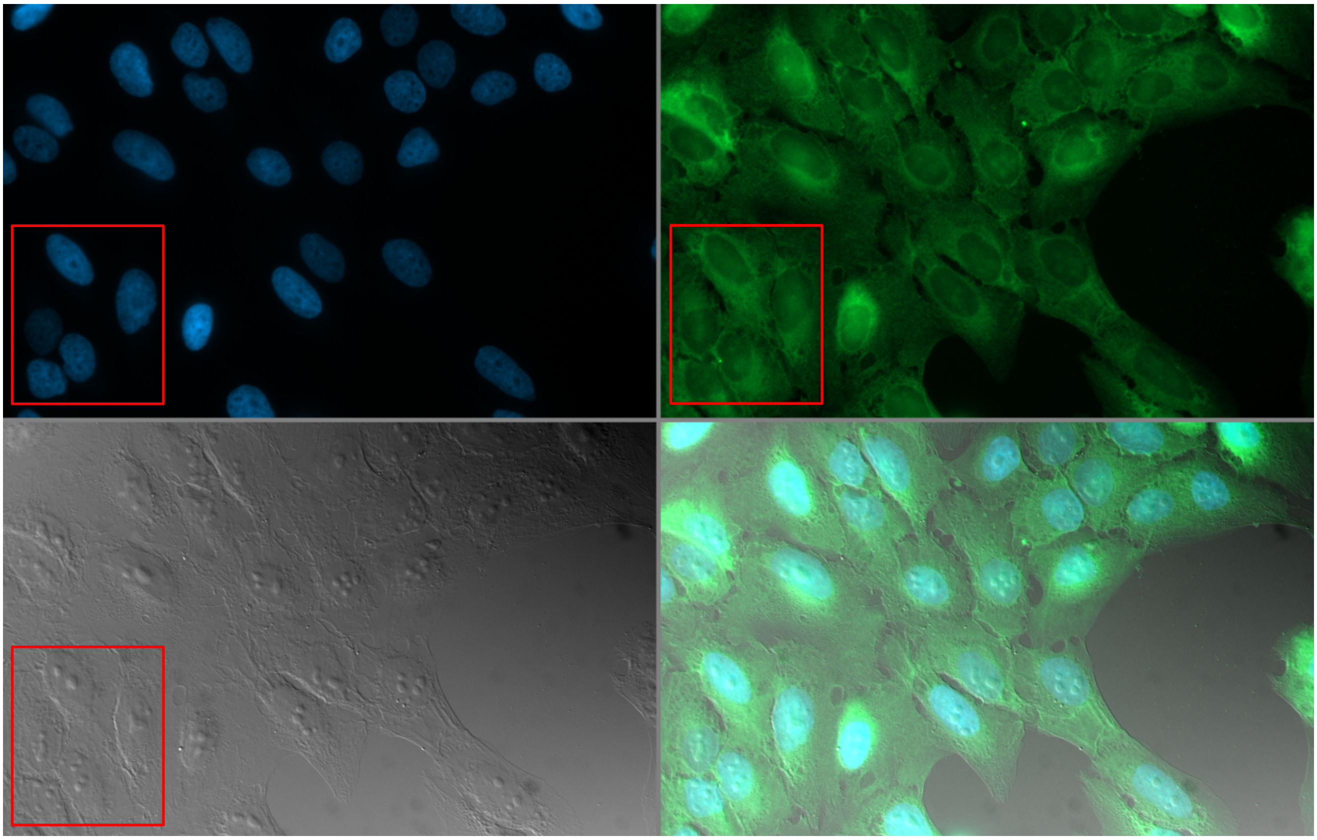

DIC Merge

**D** (relates to suppl. Fig. 1F)

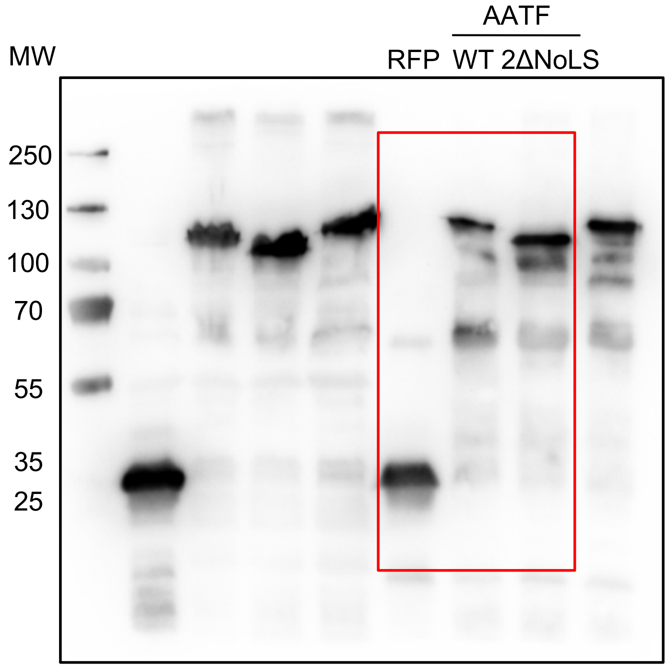

**E** (relates to Fig. 4B)

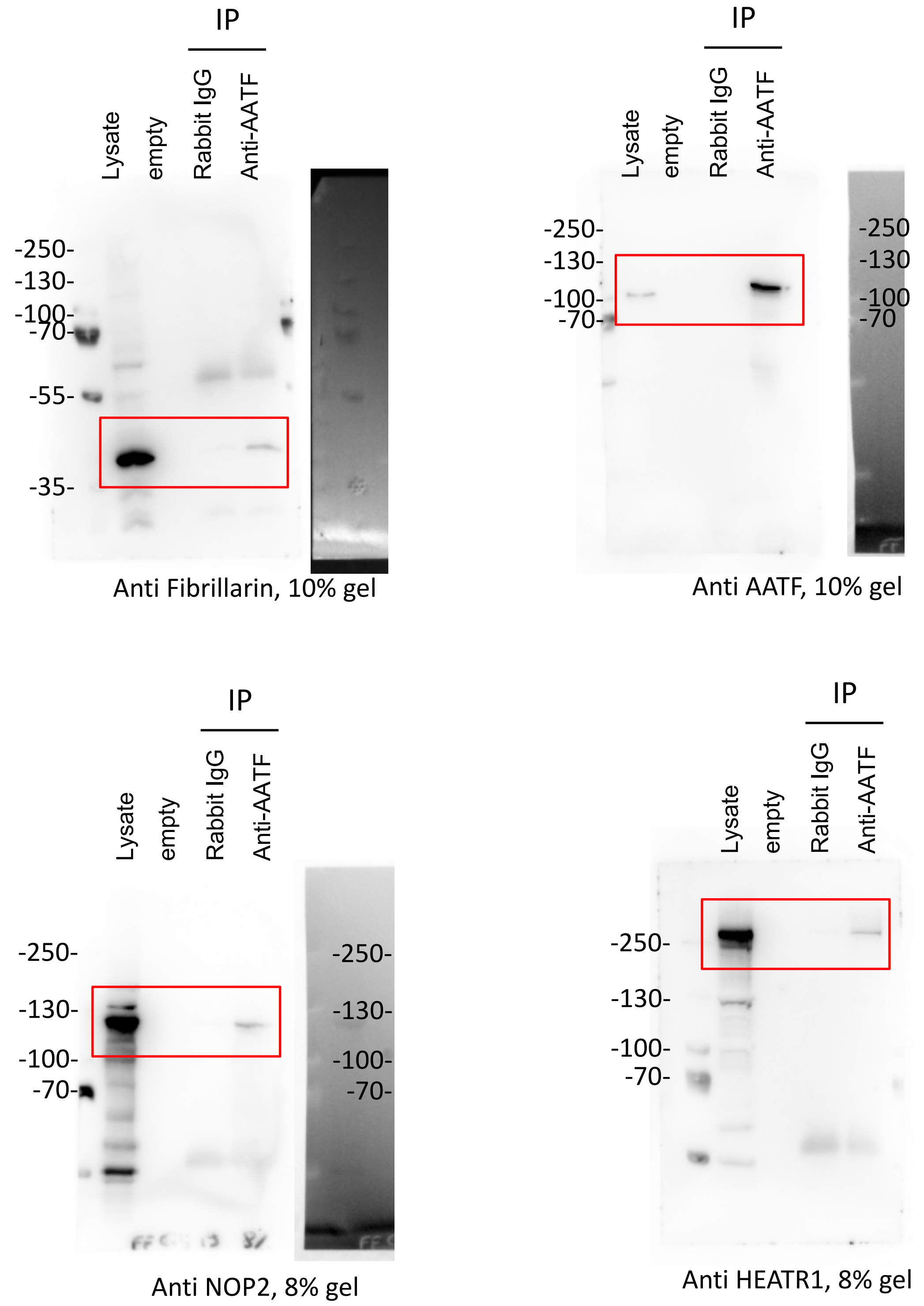
